# Systematic reconstruction of molecular pathway signatures using scalable single-cell perturbation screens

**DOI:** 10.1101/2024.01.29.576933

**Authors:** Longda Jiang, Carol Dalgarno, Efthymia Papalexi, Isabella Mascio, Hans-Hermann Wessels, Huiyoung Yun, Nika Iremadze, Gila Lithwick-Yanai, Doron Lipson, Rahul Satija

**Author notes:** These authors contributed equally (Longda Jiang, Carol Dalgarno).

## Abstract

Recent advancements in functional genomics have provided an unprecedented ability to measure diverse molecular modalities, but learning causal regulatory relationships from observational data remains challenging. Here, we leverage pooled genetic screens and single cell sequencing (i.e. Perturb-seq) to systematically identify the targets of signaling regulators in diverse biological contexts. We demonstrate how Perturb-seq is compatible with recent and commercially available advances in combinatorial indexing and next-generation sequencing, and perform more than 1,500 perturbations split across six cell lines and five biological signaling contexts. We introduce an improved computational framework (Mixscale) to address cellular variation in perturbation efficiency, alongside optimized statistical methods to learn differentially expressed gene lists and conserved molecular signatures. Finally, we demonstrate how our Perturb-seq derived gene lists can be used to precisely infer changes in signaling pathway activation for in-vivo and in-situ samples. Our work enhances our understanding of signaling regulators and their targets, and lays a computational framework towards the data-driven inference of an ‘atlas’ of perturbation signatures.

## INTRODUCTION

Recent years have witnessed an explosion in the development of new technologies for functional genomics, including the ability to measure diverse molecular modalities that span the central dogma^1–8^. The broad application of these techniques has yielded a wealth of information associating changes in molecular state across individuals, environmental conditions, and disease states^9–14^. A key challenge for the next stage of genomic analysis is to move beyond associative and correlative findings towards a more causal understanding of biological systems.

Genome engineering tools such as CRISPR hold enormous promise towards identifying causal regulators that drive molecular and functional phenotypes^15–19^, particularly when applied as part of massively parallel screens. When combined with a single-cell RNA-seq readout (i.e. Perturb-seq)^20–22^, these technologies combine the observational but unsupervised nature of RNA-seq measurements with inherent causal inference enabled by genetic perturbations. As a result, technologies like Perturb-seq offer enormous promise for causal gene regulatory network reconstruction^21–24^. For example, the Genome-Wide Perturb-seq (GWPS) resource knocked down approximately 10,000 genes in resting human cells in order to create a large-scale genotype-phenotype map and to perform an in-depth dissection of gene function^25^. This represents a powerful tool for elucidating transcriptional signatures, but applications in resting cells may fail to accurately describe context-dependent gene function.

The genomics community places significant emphasis on identifying the downstream effectors of signaling regulators to quantify and compare levels of pathway activation across diverse samples. Comparative analysis workflows routinely test for gene set enrichment using pathway signature databases, which are often compiled from diverse data types and studies^26–29^. We propose that large scale single-cell perturbation screens represent an innovative approach for refining these databases, and in particular, offer a comprehensive and data-driven workflow to enumerate transcriptional signatures and to link them to causal regulators. However, profiling dynamic gene function requires the Perturb-seq assay to be repeatedly run across multiple biological contexts, creating challenges for cost and scalability.

Here, we introduce a highly scalable Perturb-seq workflow and apply it to perturb signaling regulators across 30 separate biological contexts. We demonstrate how Perturb-seq is compatible with recent and commercially available advances in combinatorial indexing as well as a new sequencing-by-synthesis platform, sequencing 2.6 million cells in total. We divide these profiles across six cell lines and five signaling pathways in order to explore both conserved and cell type-specific responses to each signaling regulator. We introduce an improved computational framework (Mixscale) to address cellular variation in perturbation efficiency that is inherent to Perturb-seq experiments, alongside optimized statistical methods to learn differentially expressed gene (DEG) lists and broader signaling programs. Finally, we demonstrate how our data-driven inference of molecular signatures can be used to precisely infer changes in signaling pathway activation for in-vivo and even in-situ samples, including for immune and intestinal disorders. Our work enhances our understanding of signaling regulators and their targets and lays a computational framework towards the systematic construction of an ‘atlas’ of perturbation signatures.

## RESULTS

### Scalable and flexible Perturb-seq across cell lines and conditions

We aimed to use Perturb-seq to build a database of molecular response signatures, focusing on targets of different signaling regulators, and to examine how these signatures change across various cell lines. Obtaining these datasets required performing scalable Perturb-seq in a large number of different contexts, which required us to address two experimental challenges. First, we needed to create a diverse set of biological samples reflecting a diversity of cellular and environmental contexts. Second, we needed to develop a workflow to perform massively scalable Perturb-seq experiments across these multiple sample types.

To study pathway activity across multiple biological contexts, we aimed to perform Perturb-seq experiments in six different cancer cell lines from different tissues of origin: A549 (lung), MCF7 (breast), HT29 (colon), HAP1 (bone marrow), BxPC3 (pancreas), and K562 (bone marrow). To facilitate multiplexed gene knockdown screens, we modified each of these lines to express a CRISPR interference (CRISPRi) dCas9-KRAB-MeCP2 cassette^30,31^ (Supplementary Methods). To explore different environmental signaling contexts, we exposed each cell line to five distinct stimuli representing well-established pathway regulators, each of which has been broadly implicated in cellular responses and disease pathogenesis. Our selected pathways included interferon-beta (IFNβ), interferon-gamma (IFNγ), transforming growth factor beta (TGFβ), tumor necrosis factor-alpha (TNFα), and insulin (INS). Together, our combination of six cell lines and five stimuli created a diverse matrix of biological samples, which we used as input to Perturb-seq (Figure 1a).

**Figure 1.**
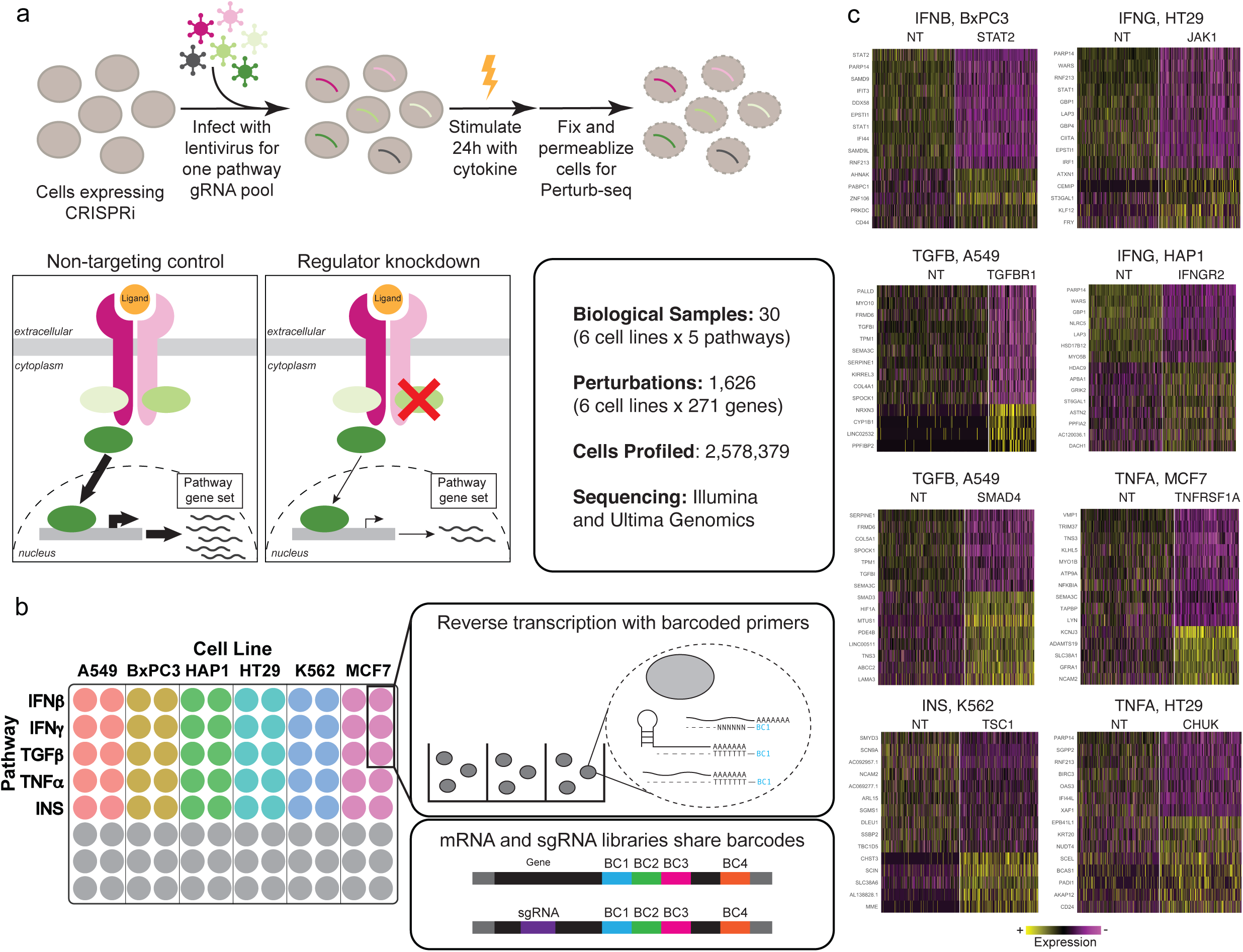
**(a).** (Top) Experimental workflow for perturbing and stimulating each pathway. (Bottom, left) Schematic showing how Perturb-seq can be used to identify pathway-specific target gene sets. (Bottom, right) Totals for biological samples, perturbations, and cells profiled in this study. **(b).** Diagram of transcriptome and sgRNA barcoding using Parse Biosciences combinatorial indexing. RNA transcripts and sgRNAs are captured, reverse transcribed, and barcoded by poly(dT) and random hexamer primers so that both modalities share a set of cell barcodes. **(c).** Eight example single-cell heatmap of one or two perturbations per signaling pathway, showing top up and down regulated differentially expressed genes.

Our goal was not to discover new pathway regulators, but instead, to characterize the molecular responses after the perturbation of known regulators. Therefore, for each pathway, we selected 44 to 61 genes based on a literature review of known regulators^32–40^. For each gene, we selected three independent single guide RNAs (sgRNA) from the Dolcetto genome-wide CRISPRi library^41^, as well as 14 non-targeting (NT) controls (Supplementary Table 1). We next created five pooled sgRNA libraries, one independently created for each pathway. For each pathway, we separately infected all six cell lines with the pathway-specific sgRNA library, and then stimulated infected cells with the corresponding cytokine for 24 hours to activate signaling (Figure 1a). This process resulted in a total of 30 distinct Perturb-seq samples.

To address logistical and scalability challenges, we adapted the Perturb-seq workflow to be compatible with the Parse Biosciences Evercode™ Whole Transcriptome Mega kit (Figure 1b, Supplementary Figures 1,2, Supplementary Methods). The Evercode kit is compatible with fixed samples, enabling us to perturb, stimulate, and fix cells with our five distinct pathway perturbation libraries at different times, and subsequently perform scRNA-seq simultaneously on all samples. This workflow also leverages three levels of combinatorial indexing to increase the scalability and cost-effectiveness of large-scale analysis^42^. Briefly, our modifications included the addition of a guide-specific primer to the cDNA amplification reaction, and the modification of the PCR reaction conditions to optimize guide recovery without adversely affecting the whole transcriptome amplification (Supplementary Figure 1,2).

We sequenced our transcriptomic libraries using a recently developed mostly natural sequencing-by-synthesis technology (Ultima Genomics)^43^, further validating a subset of samples with additional Illumina NovaSeq sequencing (Supplementary Methods). In total, we applied our combined workflow to sequence approximately 2.6 million cells across two experimental replicates. We used the set of combinatorial Parse barcodes to infer the type and the stimulation condition for each cell, and the gRNA barcode to infer its genetic perturbation. For each biological context and perturbation, our dataset enables the identification of up-regulated and down-regulated transcriptional signatures (eight representative examples in Figure 1c).

### Weighted differential expression analysis for CRISPRi Perturb-seq data

We next developed a tailored computational pipeline to optimally process and interpret our Perturb-seq datasets, aiming to solve the following problems. First, we aimed to optimize the process of identifying DEGs resulting from each perturbation, which can be challenging due to technical and biological variation in scRNA-seq assays including Perturb-seq^44,45^. Second, while our dataset encompasses thousands of individual perturbations across biological contexts, grouping these results into coherent and reproducible gene programs represents a key challenge for data interpretation. Finally, we aimed to demonstrate how to leverage these programs to interpret in-vivo scRNA-seq datasets.

We and others have previously highlighted that Perturb-seq data exhibits extensive cellular heterogeneity, even for cells that receive the same gRNA^31,46–48^. This variation encompasses both technical and biological sources, including variability in the effectiveness of perturbation. Previously, we developed a binary classification method (called ‘Mixscape’) to identify and remove cells that ‘escape’ perturbation in standard CRISPR-Cas9 knock-out experiments^46^. However, when examining well-characterized regulators in our CRISPRi dataset, we found that cells exhibited a quantitative gradient of responses (Figure 2a), consistent with multiple factors that influence variable efficiency of CRISPRi knockdown^49,50^. In these cases, binary classification is an oversimplification that can lead to the incorrect classification of ‘non-perturbed’ cells. Moreover, the existing mixscape framework assumes the presence of only a single cell type, which does not reflect our multi-context experimental study.

**Figure 2.**
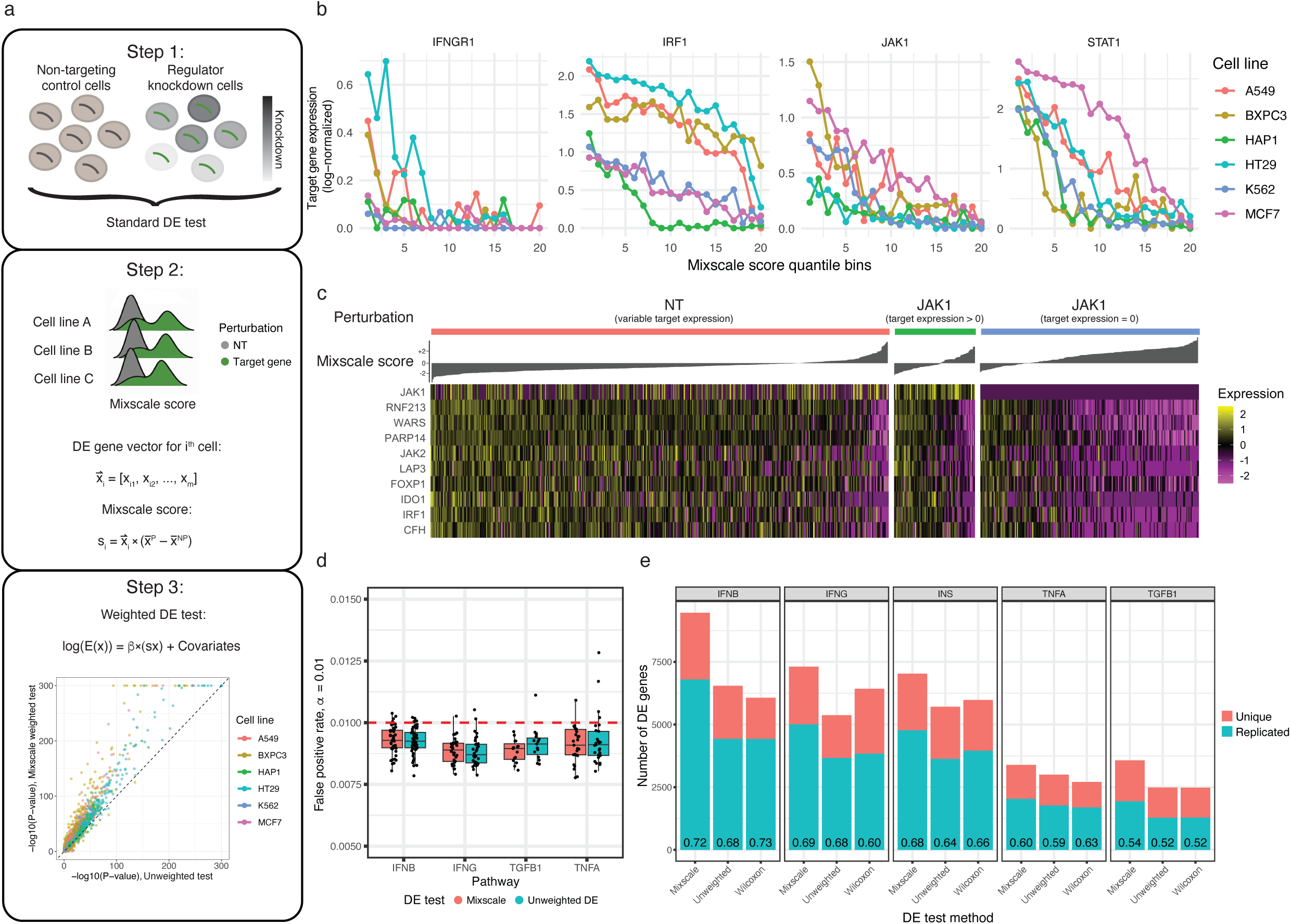
**(a).** Overview of the Mixscale scoring calculation procedure and its application in the weighted differential expression (DE) test. **(b).** Relationship between the expression levels of the perturbation targets (y-axis) and the inferred perturbation scores (x-axis) across individual cells. Expression is calculated based on pseudobulk expression of 20 bins, which cells are placed into after ordering by Mixscale score **(c).** Single-cell heatmap for JAK1 perturbation in A549 cells. Cells are ordered by Mixscale score. Even in cells where the JAK1 transcript is not detected, the Mixscale score correlates with the effective perturbation strength. **(d).** Comparison of false positive rates for the Mixscale weighted DE method (wmvReg) and the unweighted DE test computed under null simulations (Supplementary Methods). **(e)**. Replication rates of identified DE genes across scRNA-seq replicates, when applying different DE methods.

To address this, we extended the original Mixscape classifier^46^ to replace binarized classification with a continuous scalar value that reflects the strength of perturbation. Our improved approach, Mixscale, uses an initial estimate of DEG (based on a standard single cell differential expression test) from gRNA-classified cells to estimate a cell’s response (Figure 2a). As in our previous work^46^, Mixscale first estimates a cell-specific ‘perturbation vector’, representing a difference in expression between a perturbed cell and the most similar non-targeting control cells in the dataset. Instead of using this vector to infer a binarized response, Mixscale performs a scalar projection of each cell’s expression profile onto this perturbation vector, in order to quantitatively estimate each cell’s degree of perturbation (Supplementary Methods). We adapted the procedure to estimate the initial set of DEG when considering all cell lines simultaneously (to increase power), but calculated scalar projections independently for each cell line in order to robustly model potentially heterogeneous perturbation responses across biological contexts (Supplementary Methods). The output of this procedure represents a quantitative perturbation ‘score’ for each gRNA-receiving cell.

We reasoned that our perturbation score for each cell should also reflect the degree of CRISPRi-driven knockdown of the regulator itself. While the level of a single target gene is difficult to quantify accurately in individual cells, we observed a gradual decline in target gene expression with increasing perturbation scores in cells that received the target gRNA (perturbed cells) (Figure 2b). Even in cells with no detectable expression of the CRISPRi target gene, we observed a quantitative gradient of perturbation scores that revealed gradual variation in the expression of downstream regulated genes (Figure 2c). We observed similar patterns even when restricting analysis to cells receiving the same gRNA (Supplementary Figure 3a,b), indicating such variation is not purely driven by the different efficiencies among gRNAs. We replicated our findings in an independent CRISPRi Perturb-seq dataset targeting essential genes^51^. In this case, when we observed heterogeneity in perturbation scores across multiple gRNAs targeting the same gene, our molecularly-derived scores exhibited strong correlations with functional gRNA activity scores, as measured in independent proliferation-based screens from the same dataset^51^ (Supplementary Figure 3c,d). We conclude that our inferred perturbation scores represent a robust single-cell quantification of perturbation strength.

Accounting for the degree of perturbation each cell received can improve the robustness of downstream analysis. In particular, cells that exhibit weaker perturbation effects, likely driven by a lesser degree of target gene knockdown, should contribute less to DEG identification. We therefore implemented in Mixscale a weighted multivariate regression (wmvReg) procedure that accounts for multiple variables, including a cell’s perturbation score, cell line identity, and sequencing depth of each cell, in order to identify genes whose expression is dependent on gRNA identity (Supplementary Methods). The use of wmvReg to identify a final DEG list represents an iterative procedure, since it leverages cell-specific weights which themselves were constructed from an unweighted standard test. We included a ‘leave-one-out’ procedure to address potential circularity resulting from this approach (Supplementary Methods), and found that this approach mitigated artificial correlations that could otherwise arise from inaccurate initial DEG estimation (Figure 2d, Supplementary Figure 4a-c).

We further tested the statistical power of Mixscale’s wmvReg on our dataset, and compared it to alternative strategies for DEG identification. Our weighted framework identified substantially more DEG per perturbation when compared to a Wilcoxon rank sum test (Supplementary Methods) on gRNA-derived labels (on average, 404 vs. 290 DEGs per perturbation). Importantly, when we repeated the entire process on both scRNA-seq replicates separately, the DEG identified by Mixscale exhibited higher rates of reproducibility (Figure 2e, Supplementary Figure 4d), even when considering genes that were exclusively found by our procedure (Supplementary Figure 4e,f). Taken together, we conclude that Mixscale represents a sensitive, robust, and reproducible procedure to identify DEG from large scale and complex CRISPRi Perturb-seq datasets.

As we sequenced a subset of our libraries with both the Illumina NovaSeq and the Ultima UG100 platforms (Supplementary Methods), we analyzed each sequencing run independently and compared results. Gene expression estimates between the two technologies were highly correlated, with the exception of a small number of outlier genes (Supplementary Figure 5a, Supplementary Table 2), as has been previously described^52^. These platform-dependent differences in baseline expression, however, were no longer apparent after comparing NT and perturbed cells sequenced within the same platform (Supplementary Figure 5b,c). Our findings demonstrate that alternative sequencing technologies are compatible with combinatorial indexing workflows, and consistent with previous work ^25^, can be used for large-scale scRNA-seq and Perturb-seq studies.

### Conserved and context-specific perturbation responses

Our entire dataset encompasses 1,626 multiplexed perturbation experiments, where each experiment corresponds to the knockdown of a given regulator (*n* ranges from 44 to 61 depending on the pathway), in a given cell line (*n*=6), under a given pathway stimulation (*n*=5). Of these, we discarded 30 perturbations for which Mixscale identified < 5 significant DEG across cell lines. Our remaining 1,596 gene lists provide an opportunity to ask two key questions: First, do multiple regulators within the same pathway target overlapping or distinct lists of downstream genes? Second, How does a regulator’s downstream targets change across different cell lines? More broadly, our dataset highlights the necessity of learning conserved gene modules that are repeatedly identified as differentially expressed across regulators and cell lines, in order to facilitate interpretation and exploration.

Initial analysis of our data revealed both conserved and context-specific perturbation responses (Figure 3a-d). For example, key upstream regulators of the IFNγ pathway, including the IFNGR1/2 receptors and the JAK1/2 kinases, all targeted highly overlapping groups of genes. Perturbation of IRF1 resulted in only a subset of these changes (for example, STAT1 levels are unchanged after IRF1 perturbation), consistent with IRF1’s role downstream of STAT1 in interferon signaling^53^. In addition, for both the IFNγ and IFNβ pathways, we observed extensive conservation of each regulator’s downstream target genes across all six cell lines (Figure 3a,b). In contrast, we observed clear cell-type specificity in the response to TGFβ and insulin signaling (Figure 3c,d). In particular, we found that individual regulators had substantial perturbation effects in all cell lines (i.e. TSC1/TSC2), while other regulators (i.e. IRS1/GRB2) exhibited highly cell type-specific effects (Figure 3d).

**Figure 3.**
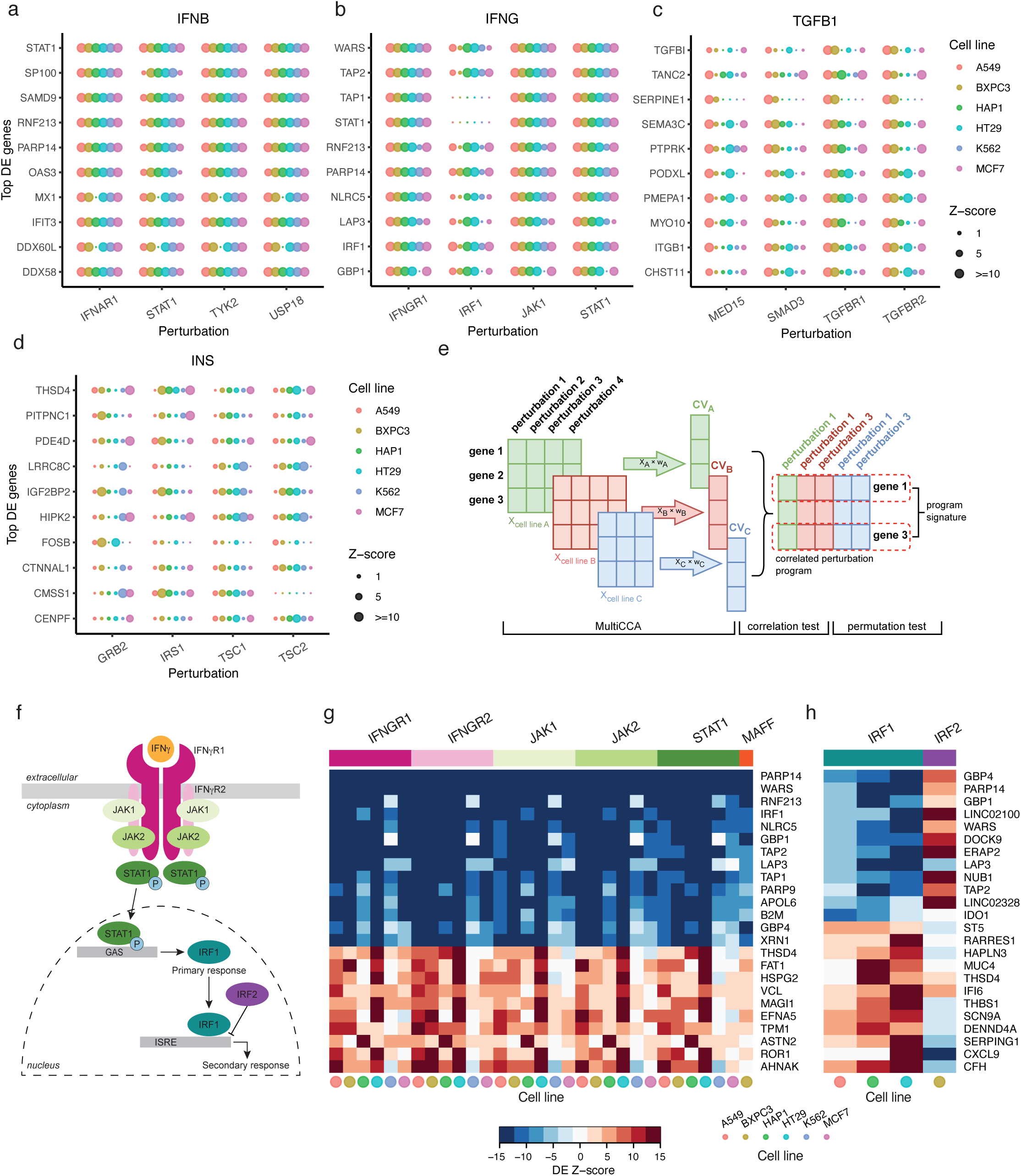
**(a-d).** Comparison of z-scores for top differentially expression genes (DEGs) (y-axis) across different perturbations (x-axis) and across six cell lines. Each dot represents a unique combination of a perturbation, a cell line, and a DEG. The size of the dot represents the magnitude of its z-score produced by the Mixscale weighted DE test. **(e).** Overview of the MultiCCA decomposition method that extracts correlated perturbations within and across cell lines (Supplementary Methods) **(f).** Overview of the main regulators in the IFNγ pathway. **(g, h).** The first two perturbation programs for the IFNγ pathway, returned by MultiCCA decomposition. Each column indicates a combination of either a positive or negative regulator (upper labels) and a cell line (bottom labels), and each row indicates a top DEG from the program signature gene list. Additional perturbation programs are shown in Supplementary Figure 6.

While the presence of 1,596 molecular signatures reflects the richness of our dataset, we sought to define consistent patterns across perturbations and cell lines, reasoning that these conserved signatures would increase robustness and interpretability. To identify shared DEG patterns across perturbations and cell lines, we adapted the MultiCCA-based^54^ decomposition approach DIALOGUE ^55^ into a pseudo-bulk-level decomposition method (Supplementary Methods). Briefly, MultiCCA identifies linear combinations of features (i.e. perturbation responses) that are highly correlated across different matrices (i.e. Perturb-seq experiments) (Figure 3e). We applied this approach separately to each of our five pathways, learning responses that we deem ‘perturbation programs’.

Importantly, MultiCCA can return multiple perturbation programs for each pathway, which can depict the hierarchical relationship shared by different regulators across cell lines. For example, when applied to the IFN! Perturb-seq datasets, the first perturbation program represents a set of hundreds of canonical targets of the IFN! pathway (Figure 3f), and is tightly conserved across cell lines (Figure 3g, Supplementary Table 3). Our approach also directionally links the expression of this program to specific regulators. We linked program 1 to the canonical positive IFN! upstream regulators (IFNGR1, IFNGR2, STAT1, JAK1, JAK2), confirming that this program represents a comprehensive description of the canonical IFN! signaling pathway (Figure 4f,g). We obtained similar results when applying our approach to IFNβ datasets (Supplementary Figure 6a), where we also identified negative regulators of the first canonical program (USP18), which is known to inhibit signaling via competitive binding to IFNAR2^56,57^.

**Figure 4.**
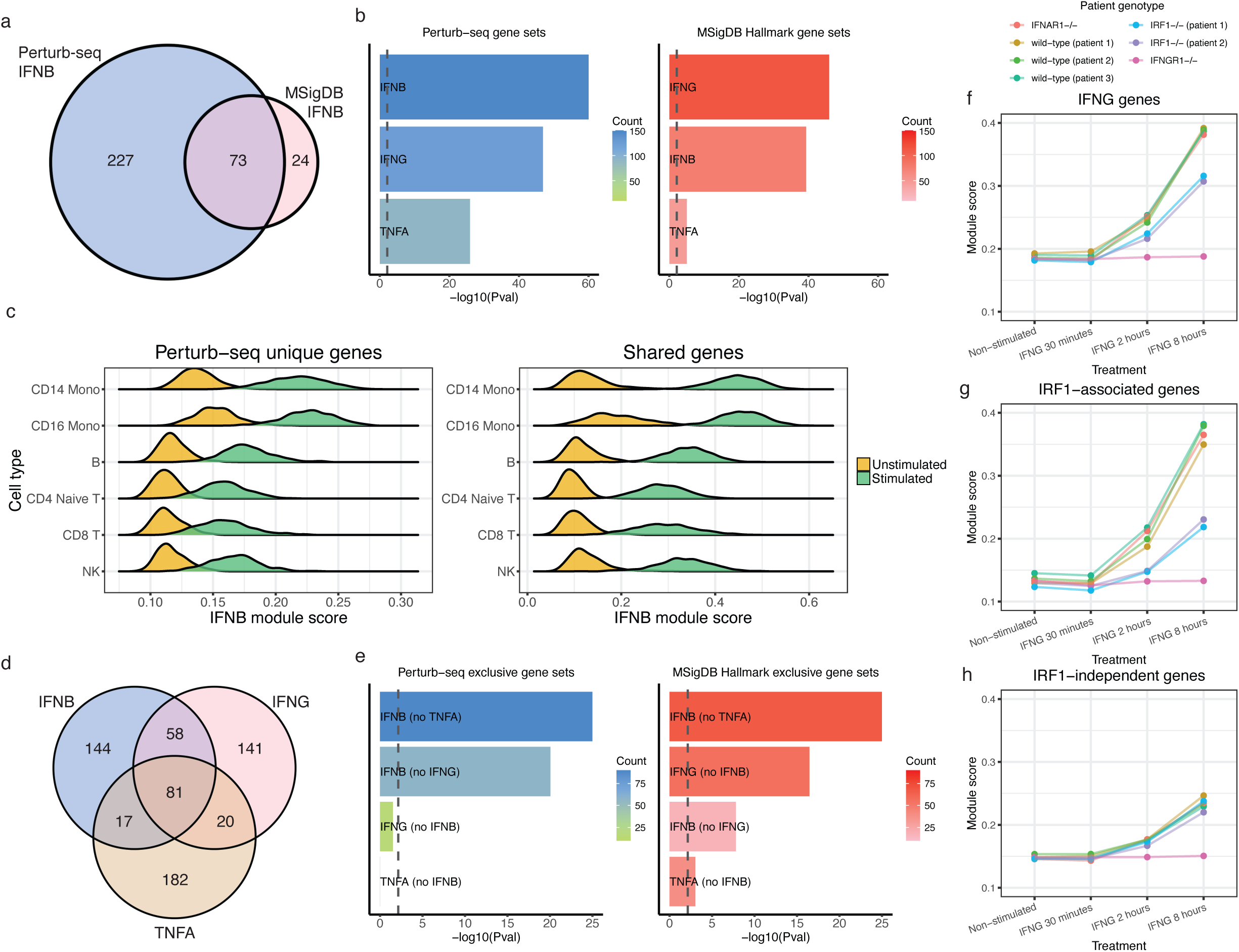
**(a).** Overlap between IFNβ program 1 genes identified by our Perturb-seq experiment and the IFNβ signatures curated by the MSigDB Hallmark collection (Supplementary Methods). **(b).** Gene set enrichments for a set of DEG from IFNβ-stimulated monocytes (Supplementary Methods), using either Perturb-seq or MSigDB signature lists. Dashed line represents the Bonferroni-corrected threshold for statistical significance. **(c).** IFNβ module score comparing unstimulated and stimulated Monocytes from an external dataset (Kang et al. 2018 Nat. Biotech) using the Perturb-seq unique gene set (the left panel) or the shared Perturb-seq and MSigDB gene set (the right panel). **(d).** Overlap of the IFNβ, IFNγ, and TNFβ pathway genes identified by our Perturb-seq experiment. **(e).** Same as (b), but for pathway-exclusive gene sets. Only the Perturb-seq gene lists correctly identify significance for IFNβ pathway lists. **(f-h).** Gene set module scores for IFNγ pathway genes, IRF1-associated genes, and IRF1-independent genes calculated in an external dataset that includes IRF1-deficient patients (Supplementary Methods). The IRF1-associated genes and IRF1-independent genes are identified using the IFNγ program 1 and 2 in our Perturb-seq data (Supplementary Methods).

The second perturbation program for IFN! specifically represents genes that are targets of a subset of more downstream regulators (Figure 3h). These include the positive regulator IRF1, which is known to act downstream of JAK/STAT signaling^53^, and to affect only a subset of pathway targets. We identified IRF2 as a negative regulator of this program, consistent with its known role in inhibiting IRF1-mediated signaling^58,59^.

Applied across our datasets, we identified 31 different perturbation programs (Supplementary Figure 6a-c, Supplementary Table 3). We note that for the INS pathway, MultiCCA failed to return clear perturbation programs due to extensive heterogeneity and minimal conservation in cell-type specific responses. To identify robust programs in this case, and to complement our cross-cell line program discovery using MultiCCA, we also used hierarchical clustering^60^ to group together shared perturbation responses for multiple perturbations *within* a given cell line (Supplementary Methods). We applied this approach to learn additional 133 gene signatures (Supplementary Table 4), which we linked to the perturbation of both positive and negative regulators within individual cell types.

### Evaluating the performance of Perturb-seq pathway signatures

We first compared our learned signaling modules with those present in MSigDB, a comprehensive resource for pathway enrichment analysis^29^. We specifically considered the “Hallmark” datasets for IFNβ, IFN!, TNFα, and TGFβ signaling. While these signatures are highly curated and leverage multiple independent publications, the underlying datasets used for creation have different origins and are collected with diverse technologies. Contrastingly, the inherently multiplexed design and uniformity of our Perturb-seq data collection minimizes batch effects and facilitates the inference of robust signatures across multiple systems. Moreover, our Perturb-seq signatures were inferred from functional genetic perturbation experiments, in contrast to observational data that is often utilized in the construction of gene signature databases^26–29^. We therefore compared our perturbation programs against each of the four hallmark gene sets.

For each pathway, we observed overlap between our learned pathway signatures and hallmark MSigDB signatures, though we did observe discrepancies as well (Figure 4a, Supplementary Figure 7a). For example, for targets of IFNβ signaling, the two databases shared 73 genes, MSigDB uniquely identified 24 genes, and Perturb-seq uniquely identified 227 genes (Figure 4a). We found that the unique Perturb-seq genes exhibited extensive reproducibility across multiple cell lines (Supplementary Figure 7b). Existing literature also suggests many of these 227 genes are genes regulated by IFNβ (e.g., BRIP1^61^, RNF213^61,62^, DDX60L^63^). Of the MSigDB-unique genes, the vast majority (70.8%, 17 out of 24) exhibited low expression levels in our Perturb-seq data, with count per million (CPM) under 20 (Supplementary Figure 7c). In comparison, only 0.4% (1 out of 227) of the Perturb-seq-unique genes had a CPM below 20. Similar observations were also made for other pathways (Supplementary Figure 7c), suggesting that the MSigDB-unique genes are either not as reliable indicators of signaling pathway activation, or cannot be accurately quantified by the Parse Evercode technology. We observed between 200 and 300 unique Perturb-seq genes for all benchmarked pathways (Supplementary Figure 7a).

We next evaluated the performance of these gene signatures to interpret experimental data from new biological contexts, but where ground truth validation of signaling pathway activation was available. For example, we considered a publicly available scRNA-seq dataset of human PBMC comparing both resting and IFN"-stimulated cells^64^. From this data, we extracted CD14+ monocytes, inferred DEG of stimulated vs. resting cells, and performed enrichment analysis against pathway signatures from Perturb-seq or MSigDB (Supplementary Methods). As expected, we observed an enrichment for IFNβ pathway genes in both signature sets (Figure 4b), with a stronger signal using the Perturb-seq signature list. To further validate the comprehensiveness of our gene list, we also performed module score analysis on both the 73 shared genes and the 227 unique Perturb-seq genes in IFNβ-stimulated CD14+ monocytes (Supplementary Methods). Our results demonstrated that both gene lists effectively distinguished stimulated cells from controls, demonstrating that the additionally identified genes from Perturb-seq analyses were reproducible hallmarks of IFNβ stimulation in a new, in-vivo biological context (Figure 4c). In comparison, the 24 MSigDB unique genes showed more limited power to strongly distinguish between stimulated and control cells, and accurate discrimination was only possible for myeloid cell types but not lymphoid cell types (Supplementary Figure 7d).

We also observed significant enrichment for alternative gene sets, including the IFNγ and TNFα pathways, using both Perturb-seq and MSigDB. This is unsurprising, as the targets of the three signaling pathways are known to overlap (Figure 4d), but illustrates the challenge in correctly inferring specific pathway activity from gene set databases. Importantly, the Perturb-seq datasets not only exhibited substantially stronger enrichment than MSigDB for IFNβ (p-value = 3.8×10^-70^ vs. p-value = 4.6×10^-40^), but also more clearly distinguished IFNβ from IFNγ signaling ( p-value = 1.3×10^-47^) (Figure 4b).

Given the improved breadth and reproducibility of our Perturb-seq gene signatures, we considered that we could leverage these data to distinguish enrichment between closely related pathways (Supplementary Methods). We repeated the enrichment test on IFNβ-stimulated cells using three sets of genes, 139 genes that were shared between Perturb-seq IFNβ and IFNγ pathways, 289 that were unique to IFNβ, and 198 that were unique to IFNγ. We found that only shared and IFNβ-exclusive gene sets exhibited enrichment, and there was no enrichment for IFNγ-exclusive targets (Figure 4e). We obtained similar results when comparing shared and unique genes for the IFNβ and TNFα pathways, which are also highly overlapping (Figure 4d). By contrast, when utilizing MSigDB signatures, exclusive gene sets for both IFNβ and IFNγ remained statistically enriched (Figure 4e).

We repeated this ground-truth validation with three additional external datasets performing IFNγ-stimulation of human PBMCs^65^, TNFα-alpha stimulation of DU145 (prostate cancer cell line)^66^, and TGFβ stimulation of OVCA420 (ovarian cancer cell line)^66^ (Supplementary Figure 8a-c). In each case, we were able to leverage our Perturb-seq datasets to correctly identify the underlying stimulation pathway, and to generate exclusive gene sets that correctly excluded enrichment for closely related pathways (Supplementary Figure 8 a-c). Notably, in all four of these ground-truth validations, the external datasets utilized biological systems that were not present in our Perturb-seq datasets.

Lastly, we asked whether our datasets could be used to correctly identify not only broad pathway programs, but to accurately identify specific sub-programs driving cellular responses. We considered a recent bulk RNA-seq dataset of mycobacteria-exposed human fibroblasts from patients with an inherited IRF1 deficiency^67^. When compared to control populations, IRF1-deficient fibroblasts exhibited deficient IFNγ signaling responses (Figure 4f). However, when considering exclusive gene sets to separate IFNγ response genes into IRF1-associated and IRF1-independent groups (Supplementary Methods), we correctly identified that only IRF1-associated genes exhibited impaired transcriptional activation, while independent genes were unaffected (Figure 4g,h). Strikingly, we were able to correctly infer IRF1-specific enrichment even in datasets from non-human species, including RNA-seq data from virally infected IRF1-deficient bat cells^68^ (Supplementary Figure 8d). We conclude that our Perturb-seq signatures outperform existing databases, and can be used to sensitively infer signaling pathway activation in diverse external datasets.

### Inferring signaling pathway activation for in-vivo and in-situ datasets

Having demonstrated the ability to correctly infer causal regulatory events from in-vitro experiments with ground-truth, we next aimed to extend these analyses to infer pathway activity in new in-vivo datasets. For example, the activation of interferon signaling has been widely reported to affect multiple cell types after SARS-CoV-2 infection^69–74^. However, both type I (IFN#/β)^69–71^ and type II (IFN!)^72–74^ are widely reported to contribute to cellular responses to infection. While it is traditionally challenging to disentangle the specific effects of these pathways, even with scRNA-seq datasets from healthy and SARS-CoV-2-infected samples, we reasoned that our Perturb-seq pathway signatures could help determine whether individual cell types were responding exclusively to type I, type II, or both interferon pathways.

To address this, we leveraged a large-scale scRNA-seq dataset from COVID-19 patients. We utilized a dataset of circulating immune cells from the COvid-19 Multi-omics Blood ATlas (COMBAT)^75^, and determined cell-type specific DEG for myeloid and lymphoid cell types (Supplementary Methods). Using the full pathway signatures from Perturb-seq, we observed strong enrichment in many cell types for both IFNβ and IFNγ targets (Figure 5a). Alongside general heterogeneity in patient responses (Figure 5b-c), we confirmed a depletion of interferon responses within critically ill patients, and localized this deficiency most strongly to T cell subgroups (Figure 5a). When searching for enrichment of exclusive gene sets, we only identified enrichment in IFNβ-specific groups, with no enrichment for IFNγ-specific groups (Figure 5a,d,e). Consistent with this result, when ordering patients by their molecular response to disease (Supplementary Methods), we observed a clear gradient of expression for IFNβ-specific gene sets, but not IFNγ-specific gene sets (Figure 5d,e). These results highlight the specific role of IFNβ in the response of circulating immune cells to SARS-CoV-2 infection.

**Figure 5.**
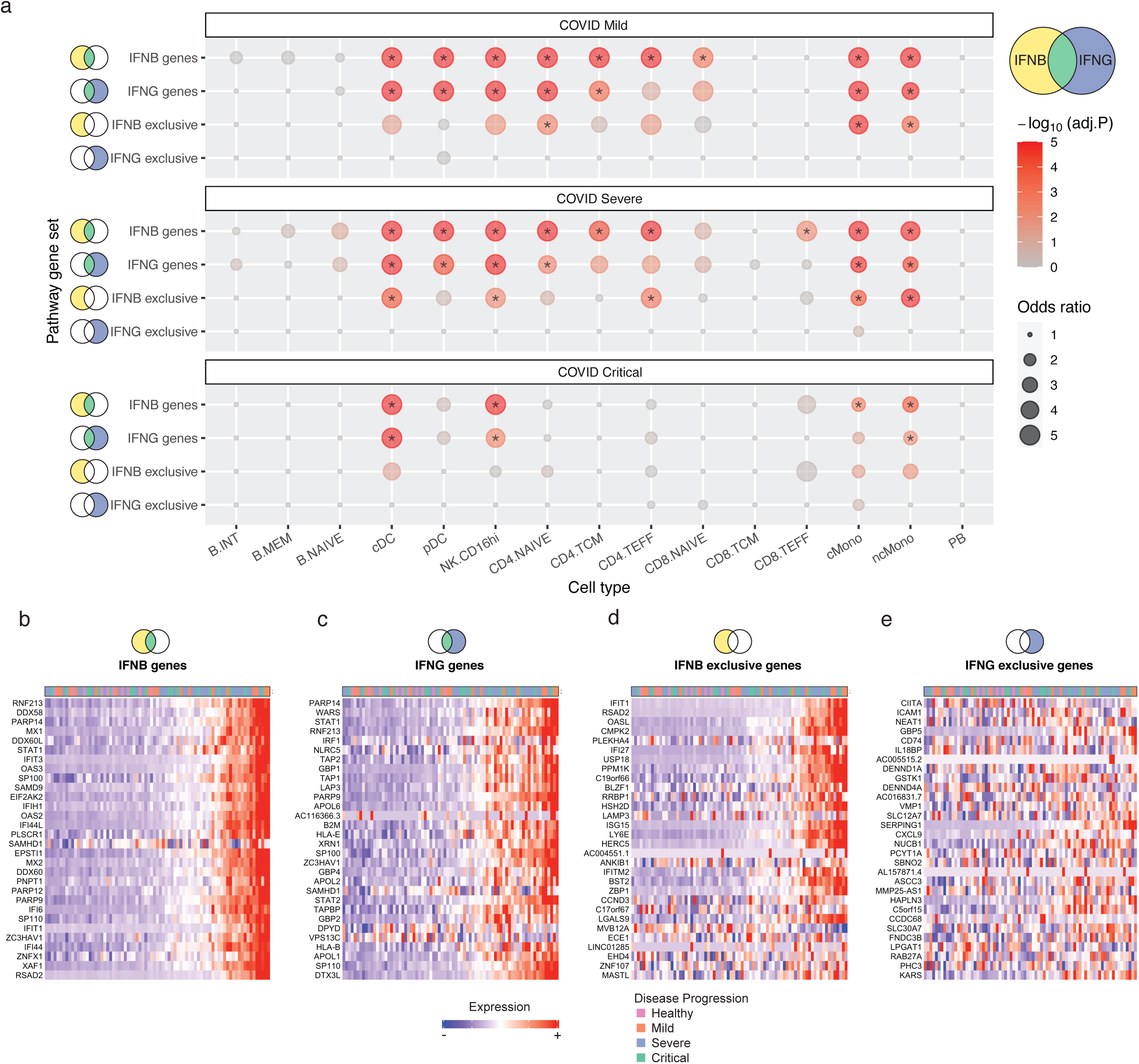
**(a).** Enrichment test for DEG across COVID-19 severity groups from an external dataset (COvid-19 Multi-omics Blood Atlas, COMBAT). Rows represent gene sets from our Perturb-seq data, columns show cell types yielding DEG between disease and healthy cells. Dot size denotes the odds ratio, color intensity indicates the adjusted P-value (Benjamini-Hochberg; * indicates p < 0.01). **(b, c).** Expression heatmap for the top 30 IFNγ and IFNβ pathway genes (including shared genes). Each column represents pseudobulk expression of CD14 monocytes within each individual (Supplementary Methods). Columns are ordered by increased expression of a combined gene list of IFNγ and IFNβ genes **(d, e).** Same as (b-c), but for pathway-exclusive gene lists. Only IFNβ-exclusive gene sets are coordinately up-regulated, consistent with the enrichment analysis in (a).

In addition to identifying disease-relevant pathways, we reasoned that our Perturb-seq datasets could also help to prioritize and pinpoint specific cell types that may be driving disease state. We first considered a comprehensive dataset of immune and epithelial cells from patients with Crohn’s disease (CD)^76^, an immune-mediated inflammation disorder known to primarily affect the gastrointestinal tract, leading to symptoms such as abdominal pain, severe diarrhea, and malnutrition^77^. Notably, anti-TNFα therapy has been repeatedly demonstrated to be an effective treatment for CD patients, but it is unclear if there are specific cell types that are responding to treatment.

The original manuscript computed DEG between healthy and CD colons for 54 cell types, identifying broad enrichment for inflammatory pathways without observing specific enrichment for TNFα signaling pathways^76^. Re-analyzing these DEG sets, we identified clear and specific up-regulation of TNFα targets, with minimal contributions from interferon-signaling pathways. TNFα-enrichment is observed primarily in non-immune subsets, including enterocytes, epithelial cells, goblet cells, and fibroblasts (Supplementary Figure 9). Interestingly, we found that only specific subgroups of these cells exhibited enriched TNFα signaling. For example, of five fibroblast subgroups, only two exhibited activation of TNFα targets. We observed similar heterogeneity in enterocytes and goblet cells, identifying specific sub-clusters with TNFα target-enriched DEG, alongside additional subclusters with no enrichment. These findings highlight specific subsets of non-immune cells that likely reflect promising cellular targets of anti-TNFα.

Finally, while our previous analyses focused on scRNA-seq datasets, gene signatures can be applied to broad genomic data types including spatial analyses. To demonstrate this, we explored recently generated 10x Visium spatial transcriptomic maps of the mouse colon, taken during the course of dextran sodium sulfate (DSS) colitis, which represents acute colonic injury and triggers a wound healing response^78^. We reasoned that we could implement our perturbation programs in order to better understand specific pathways and tissue regions that were influenced by disease pathology. For each of our Perturb-seq signaling programs, we computed a ‘signaling induction’ score for each region of the colon, based on the differential expression of signaling genes between voxels in the Day 14 (7 days of DSS administration followed by 7 days of healing) and Day 0 (healthy) samples (Supplementary Methods).

We observed a clear enrichment of our inferred TGFβ signaling program in discrete and specific regions of the mouse colon (Figure 6a-c). The region of highest activation (cluster 12) was located towards the center of the tissue, and exhibited striking overlap with a region annotated as exhibiting signs of ‘inflammation and hyperplasia’ based on a pathologist’s analysis of hematoxylin and eosin staining of the tissue section^78^ (Figure 6b). Following the original study’s methods for ‘digitally unrolling’ the colon ^78^ (Supplementary Methods), we further demonstrated that the induced signaling response was strongest at the distal end of the proximal-distal axis (Figure 6d,f). We observed minimal up-regulation in this region of other inflammatory signaling pathways (Figure 6c), or an existing literature TGFβ gene set used in the original study^78,79^ (Figure 6e). TGFβ represents a critical component of wound healing, and is particularly important for driving cellular proliferation and the generation of new connective tissue^80–82^. Consistent with this, the regions associated with activated TGFβ signaling were also identified in the original study as being enriched for proliferating epithelial stem cells, which help to coordinate the healing response^78^. We conclude that our Perturb-seq derived gene sets can be used to infer spatially restricted patterns of signaling activation.

**Figure 6.**
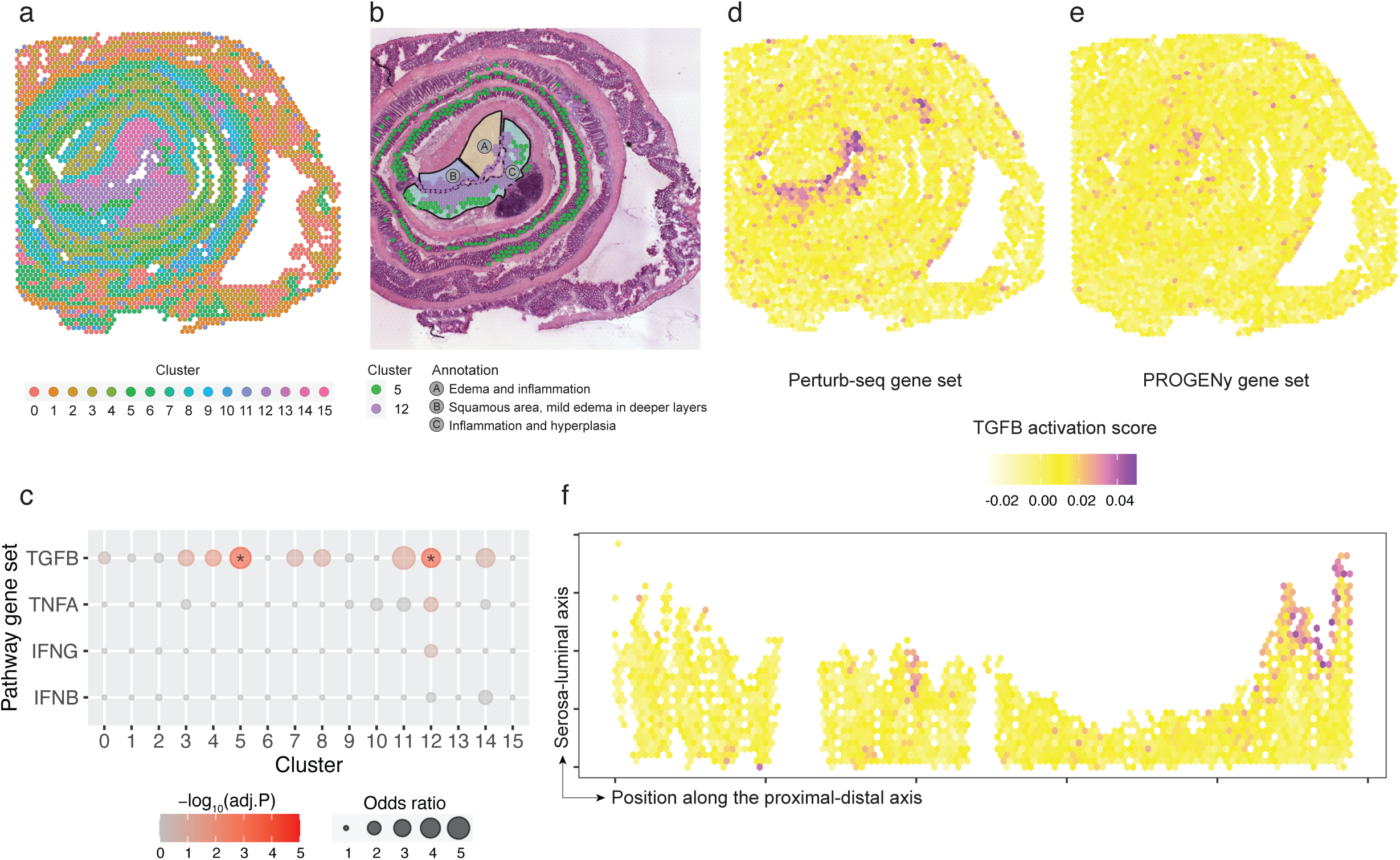
**(a).** Unsupervised transcriptomic clustering of the mouse healing intestine Visium dataset (Parigi et al. 2022 Nat. Comm.). **(b).** Overlap between the clusters with elevated TGFβ activation and the anatomical regions annotated as exhibiting signs of ‘inflammation and hyperplasia’ based on the pathologist’s analysis in the original study. **(c).** Enrichment analysis for DEG identified for different clusters in the mouse healing intestine (Supplementary Methods). Rows represent gene sets from our Perturb-seq data, columns show cell types yielding DEGs between disease and healthy cells. Dot size denotes the odds ratio, color intensity indicates the adjusted P-value (Benjamini-Hochberg; * indicates p < 0.01). **(d, e).** The TGFβ activation scores in the mouse intestine before unrolling using our Perturb-seq TGFβ gene set (d) and the PROGENy TGFβ gene set (e). **(f).** The TGFβ activation scores in the digitally unrolled mouse intestine using our Perturb-seq TGFβ gene set. The Visium spots shown in (a) are digitally flattened into a proximal to distal direction from left to right on the x-axis (Supplementary Methods).

## DISCUSSION

In this study, we adapted a commercially available combinatorial indexing workflow to perform scalable Perturb-seq experiments, and utilized this approach to learn perturbation gene expression signatures across a diverse range of signaling pathways and biological contexts. To address the challenge of heterogeneous gene expression knockdown in CRISPRi experiments, we introduce Mixscale, which infers the level of transcriptome-wide perturbation in individual cells and enables a weighted DE testing framework for boosting statistical power. We utilized these data to identify reproducible pathway (and sub-pathway) level signatures that were conserved across multiple cell types. We found that our signatures broadly extended existing and widely used gene sets, that they could be used to accurately infer signaling activitys in-vivo when examining bulk, single-cell, or spatial transcriptomic datasets.

Our study was inspired by previous efforts, including Genome-Wide Perturb-seq (GWPS)^25^, to learn large sets of gene signatures in a data-driven way. However, GWPS was mainly conducted in a single cell type, and the scalability of the study was driven by the need to profile more than 10,000 genetic perturbations. Our study does not pursue a genome-wide perturbation approach, but instead, our scalability constraints were driven by the large number of biological contexts, necessitated by the goal of learning signaling pathway signatures (which could not be observed by perturbing regulators at steady state). We addressed this challenge using the Parse Biosciences and Ultima Genomics platforms, but note that there are a series of pioneering combinatorial methods that enable larger scale Perturb-seq experiments^83,84^. In addition to the inherent multiplexing of Perturb-seq, where perturbed and non-targeting control cells are simultaneously profiled in the same experiment, performing Perturb-seq on fixed samples enabled us to multiplex different cell types and signaling pathways together, further reducing batch effects.

Our efforts are complementary to alternative approaches to learn gene expression signatures. For example, direct stimulation of cells with signaling ligands can also be used to measure transcriptome-wide signaling responses^79,85–89^. While this approach does not require additional genetic perturbations, it may also trigger the downstream activation of secondary pathways (which would be measured together with the primary targets), and cannot be used to infer sub-pathway level signatures. Notably, both approaches benefit from multiplexed single-cell profiling. For example, the Immune Cell Dictionary utilized multiplexed single-cell analysis to profile responses of immune cell types to 86 cytokines^89^. Given the rapidly growing number of identified cell types, multiplied by the vast possibilities for single or combinatorial genetic and environmental perturbations, we expect that the generation of response signature dictionaries will represent a primary use case for new massively scalable single-cell sequencing techniques.

The experimental design of this study introduces several limitations to the conclusions we can draw with our molecular pathway signatures. First, our coverage of biological systems is not comprehensive and represents only an initial set of cell types and stimulation conditions. As other groups generate additional data, these new datasets could be integrated with our own to generate improved gene signatures. Second, our experimental design collects cells after stimulation at a single time point, which prohibits the study of temporal signaling dynamics and presents another opportunity for future Perturb-seq experiments. Third, while our use of a single profiling technology minimizes batch effects, our signatures may exclude genes that are not well-quantified by Parse Biosciences, even with large numbers of cells^90^. Finally, while our primary goal in this study was to identify conserved pathway genes across different cell lines, our datasets also generated 141 cell-line specific response lists (Supplementary Table 4), enabling future studies to more deeply explore cell-type-specific responses.

In future studies, the experimental and computational framework we developed can be applied in different biological contexts and to different cellular processes to identify additional gene signatures. While our study focused on transcriptional signatures, incorporating additional modalities, such as chromatin accessibility^91–93^ and protein levels^94^, could provide further insights. We could also employ combinatorial perturbation technology^95^ to explore interactions between multiple regulators within and across pathways. All these would greatly enrich our understanding of signal transduction at multiple steps of the central dogma.

## AUTHOR CONTRIBUTIONS

L.J., C.D., E.P., and R.S. conceived the research. C.D., E.P., I.M., H.H.W., and H.Y. performed experimental work. L.J. performed the computational work and developed the software tool with guidance from R.S.. N.I., G.L.Y., and D.L. performed the Ultima sequencing and generated the simulated paired-end fastq data. All authors participated in interpretation and in writing the manuscript.

## ACKNOWLEDGEMENTS

We thank all members of the Satija Lab at New York Genome Center for useful discussion. We acknowledge the authors of the external datasets used in this study for making their valuable resources publicly available. This work was supported by the Chan Zuckerberg Initiative (EOSS5-0000000381, HCA-A-1704-01895 to R.S.), and the National Institutes of Health (RM1HG011014-02 and 1OT2OD033760-01 to R.S).

## DATA AND CODE AVAILABILITY

Processed data is available at Zenodo^96^ (https://zenodo.org/records/10520190). Software implementing our approach is freely available as an open-source R package *Mixscale* (https://github.com/longmanz/Mixscale). A vignette demonstrating the application of *Mixscale* is also available as an online resource (https://longmanz.github.io/Mixscale/).

## SUPPLEMENTARY FIGURES

**Supplementary Figure 1.**
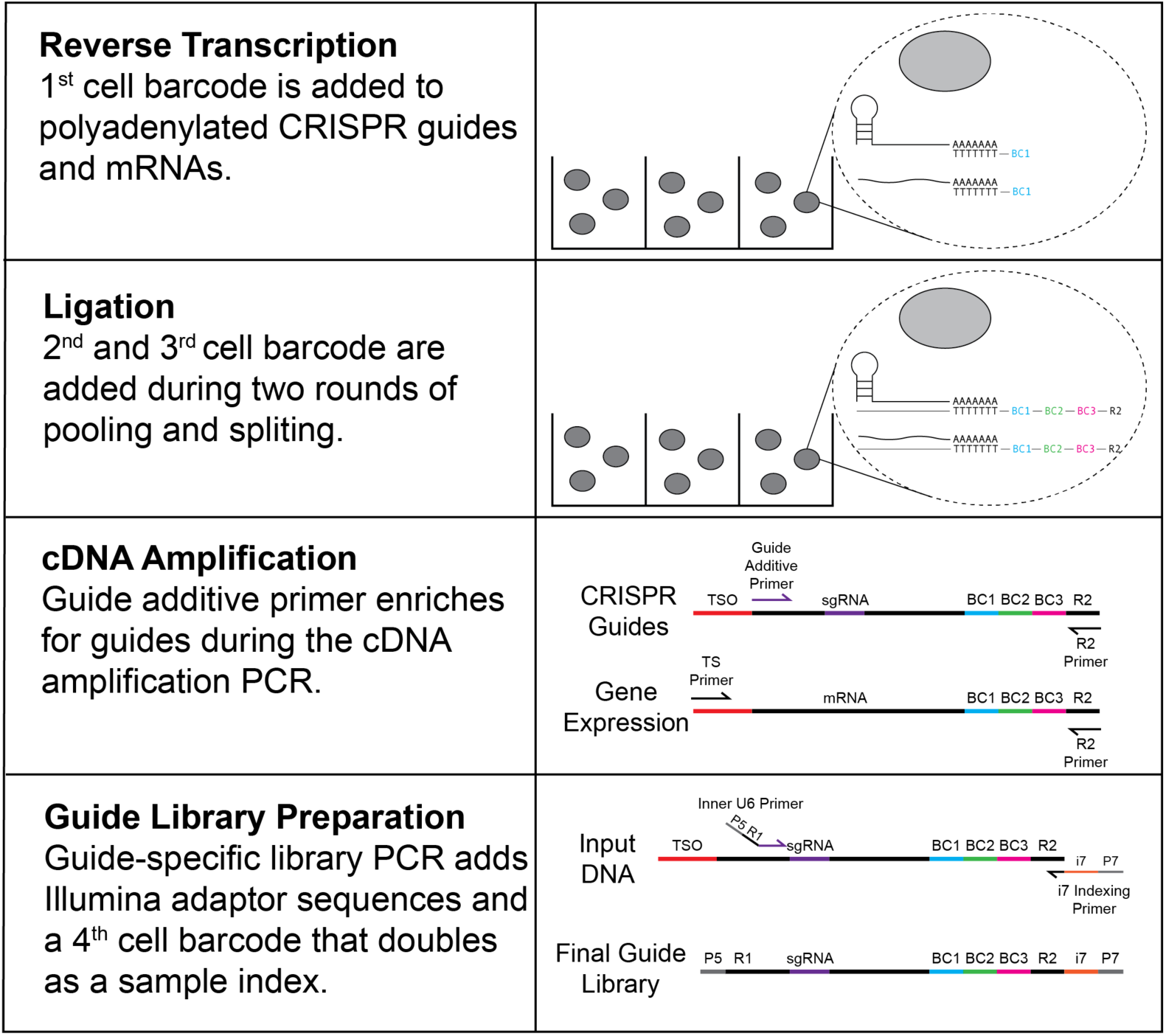
Schematic diagram of sgRNA capture, barcoding, and library preparation compatible with the Parse Bioscience Evercode^TM^ Whole Transcriptome Mega kit. Please see Supplementary File 1 for a full protocol.

**Supplementary Figure 2.**
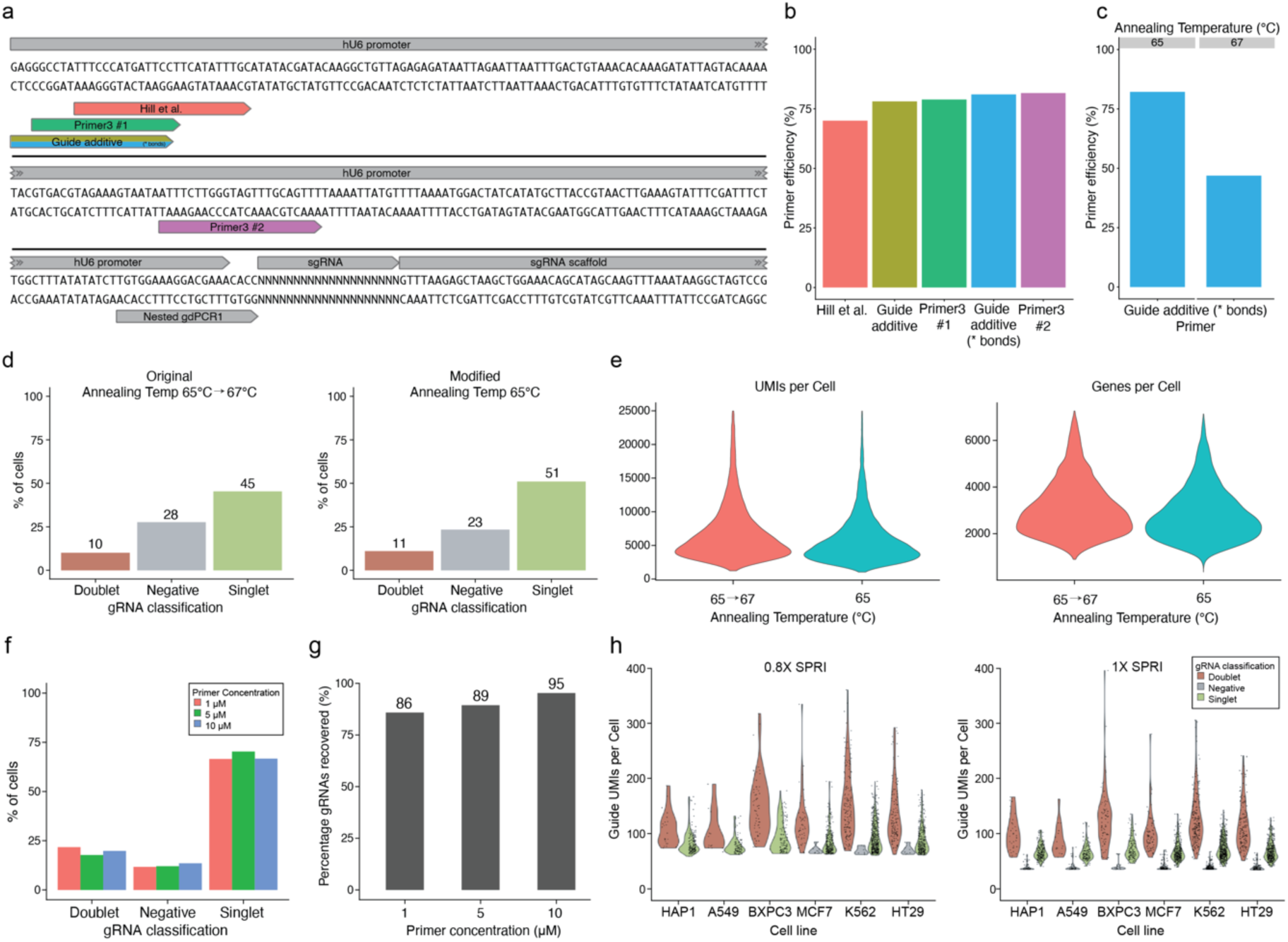
**(a).** Schematic showing the location of possible guide additive primer binding sites. **(b).** Primer efficiency for cDNA amplification, measured by qPCR with annealing temperature 65°C. Guide additive with phosphorothioate (*) bonds was ultimately chosen. **(c).** Primer efficiency of chosen guide additive primer for cDNA amplification, measured by qPCR at different annealing temperatures. **(d).** Percentage of cells assigned to each guide classification when using the original Parse cDNA amplification annealing temperature (some cycles at 67°C and some cycles at 65°C) compared to our modified annealing temperature (all cycles at 65°C). **(e).** RNA UMI counts and genes per cell when using the original Parse annealing temperature compared to our modified annealing temperature. **(f).** Percentage of cells assigned each guide classification when using different guide additive primer concentrations. **(g).** Percentage of unique gRNAs recovered when using different guide additive primer concentrations. **(h).** Guide UMIs per cell, split by guide classification, when using the original cDNA amplification SPRI bead cleanup ratio (0.8X) compared to our modified ratio (1X). We chose a 1X cleanup to help retain shorter guide transcripts captured by random hexamer reverse transcription primers.

**Supplementary Figure 3.**
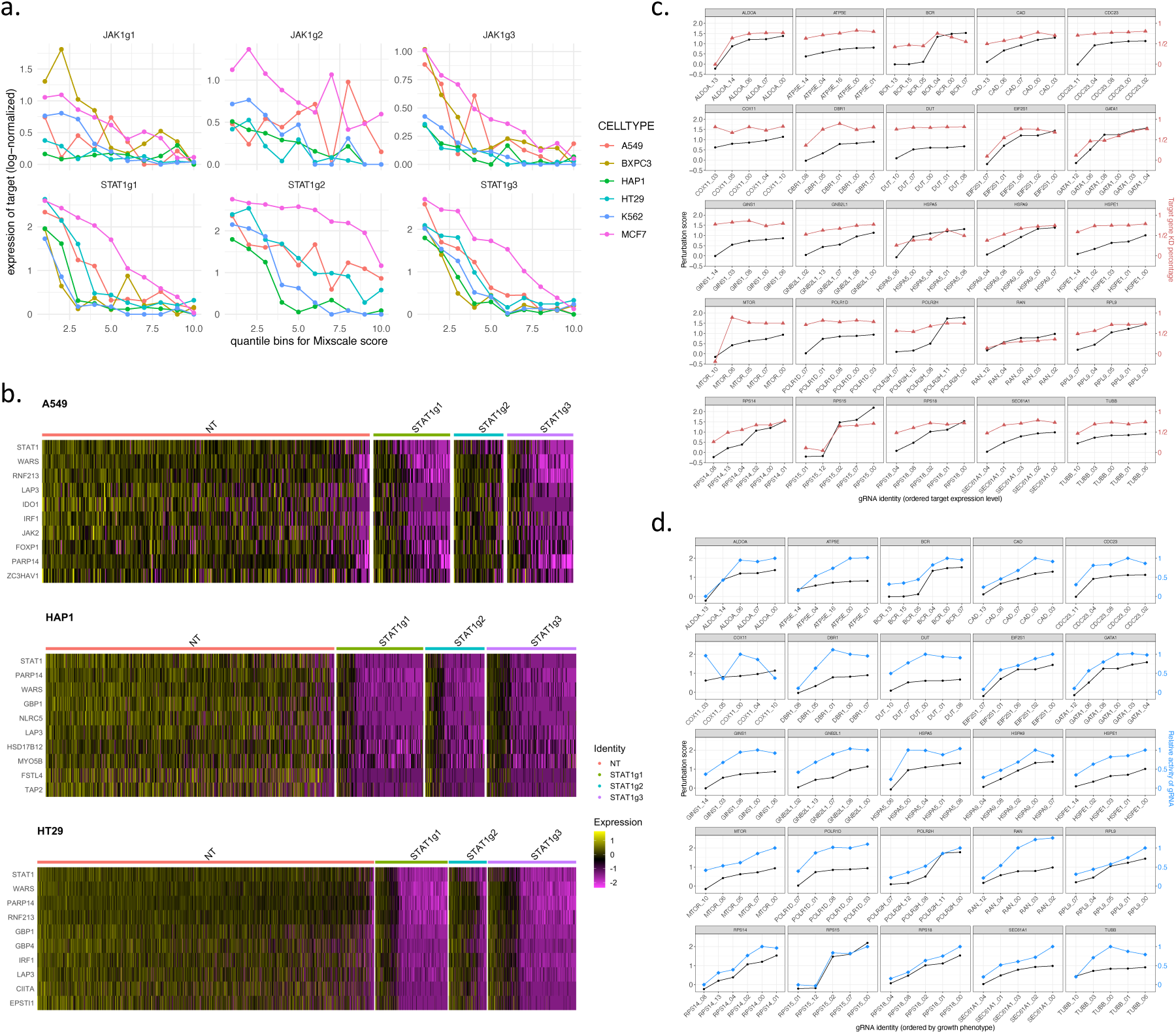
**(a)** Scatter plots illustrating the relationship between the expression level of the perturbation targets (y-axis) and the perturbation scores (x-axis) in each cell. This plot is analogous to Figure 2b but this time cells are stratified by their guide RNA identities instead. **(b)** Single-cell heatmap for STAT1 perturbation in three cell lines after IFNψ stimulation, split by gRNA identities. **(c)** Comparison of Mixscale score and target gene expression estimated in an external CRISPRi dataset (Jost et al. 2020 Nat. Biotech). The figure displays the Mixscale score (y-axis on the left) using black dots and the degree of knockdown of the target gene (y-axis on the right) marked by red triangles. The x-axis represents different gRNAs, including the perfectly matched gRNA (“_00”) and those with varying numbers of mismatched nucleotides. Plot shows that gRNA that result in more effective knockdown also result in cells with higher Mixscale scores. **(d)** Comparison of Mixscale score and relative activity of gRNAs. Similar plot as in (a), but instead the figure contrasts the Mixscale score (black dots) with the relative activity of the gRNA (y-axis on the right) marked by blue diamonds, a phenotypic measure of cellular growth defects measured from a viability screen (Jost et al., 2020, Nat. Biotech). Plot shows that gRNA with the highest phenotypic activity also yield cells with the highest Mixscale scores.

**Supplementary Figure 4.**
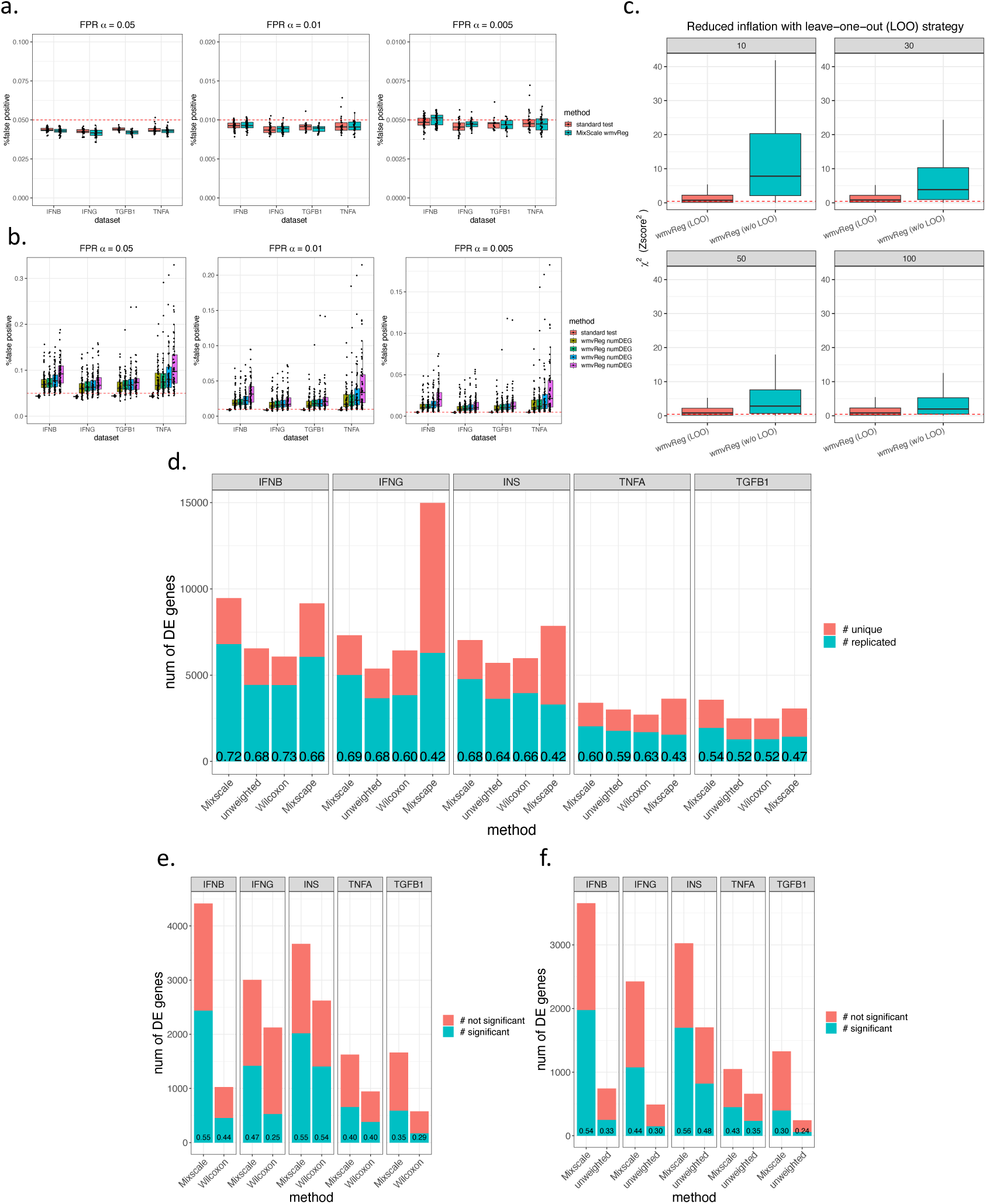
**(a).** Comparison of false positive rates (FPRs) for the Mixscale weighted DE test (wmvReg) and a standard unweighted DE test. FPRs are calculated based on alpha ≤ 0.05, 0.01, and 0.005. Genes were simulated as null by shuffling cell gRNA labels, and shuffling was performed after calculating Mixscale scores (Supplementary Methods). **(b).** FPR calculations when shuffling is performed prior to calculating Mixscale scores. This situation is a conservative control, as we force the assignment of mis-specified Mixscale scores, though in a real dataset the lack of an initially identified DEG set would abort the procedure (Supplementary Methods). **(c).** Boxplots for Mixscale DE test scores *X^2^ (= Zscore^2^*) of the genes used in the calculation of these mis-specified scores, comparing methods with and without a leave-one-out (LOO) strategy (Supplementary Methods). The red dashed line indicates the expected !^!^ under the null. **(d).** Replication rate of the DEGs across two scRNA-seq replicates using different DE methods. Same as Figure 2e but includes an approach where Wilcoxon sum rank tests are applied to Mixscape-derived labels. **(e, f).** Replication rate of DE genes that were uniquely identified by any DE method, compared to others. For example, in the IFNβ pathway dataset (Replicate 1), wmvReg and Wilcoxon uniquely identified 4,415 and 1,025 DEG, respectively, across regulators, and the plot shows the percentage that reproduce in the second replicate.

**Supplementary Figure 5.**
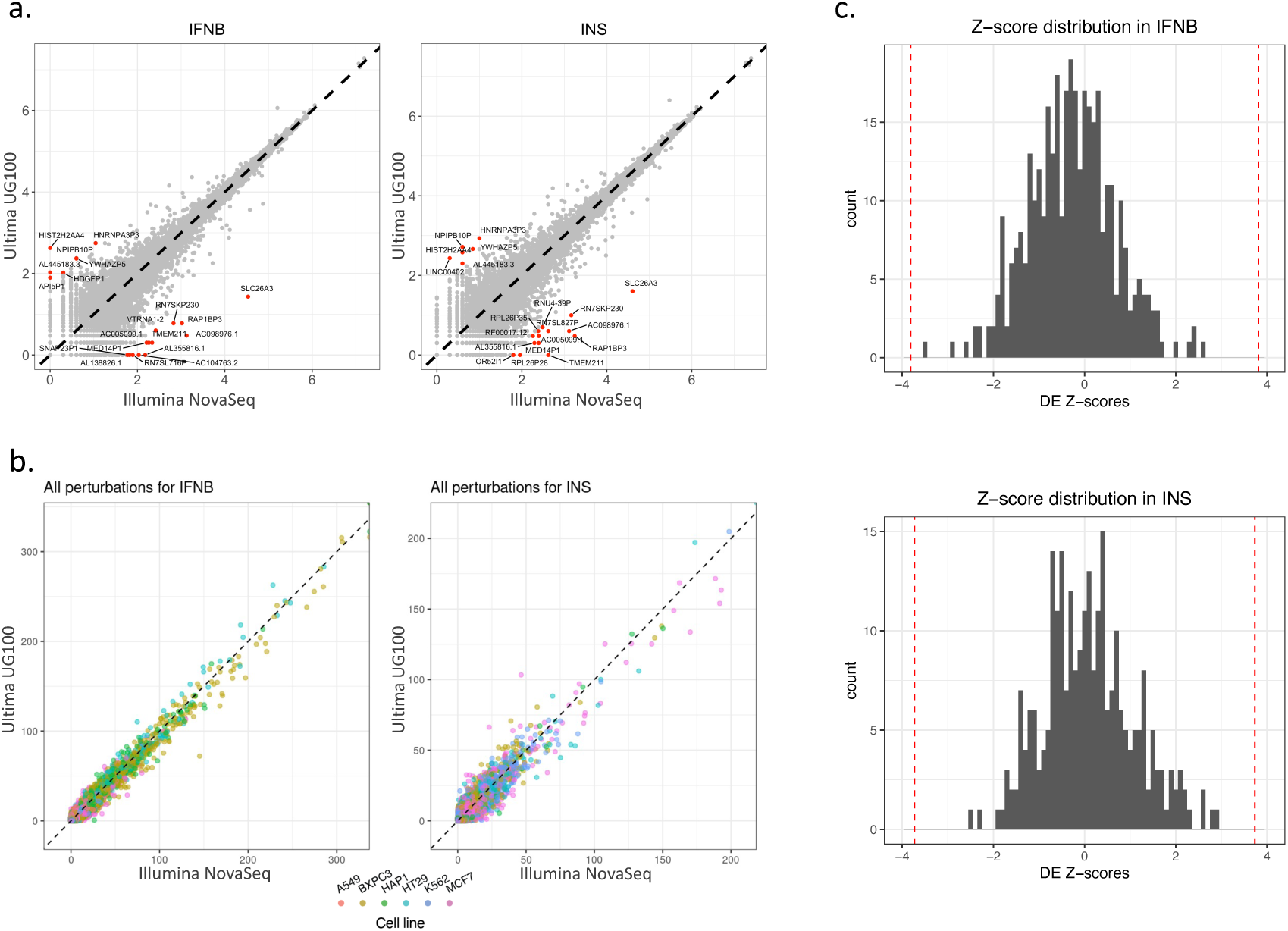
**(a).** -log_10_(Count per gene) comparison between Illumina and Ultima Genomics sequencing platforms. Data reflects pseudobulk values (all cells) for a Perturb-seq library sequenced by both Illumina and Ultima platforms, followed by the same alignment and pre-processing steps. 20 outlier genes are highlighted in red. **(b).** -log_10_(DE P-values) comparison between NT control cells and gRNA targeted cells, for the same library sequenced by either Illumina or the Ultima Genomics sequencing platform. The datasets are independently processed with Mixscale, and we observe high consistency across the sequencing platforms. **(c).** Distribution of the DE Z-scores for the outlier genes in panel (a). The red dashed lines are the Z-score threshold after Bonferroni correction for multiple testing (adjusted p-value = 0.05). Plot shows that genes that are differentially detected between Illumina and Ultima data no longer show differential expression when comparing NT and targeted cells within the same platform.

**Supplementary Figure 6.**
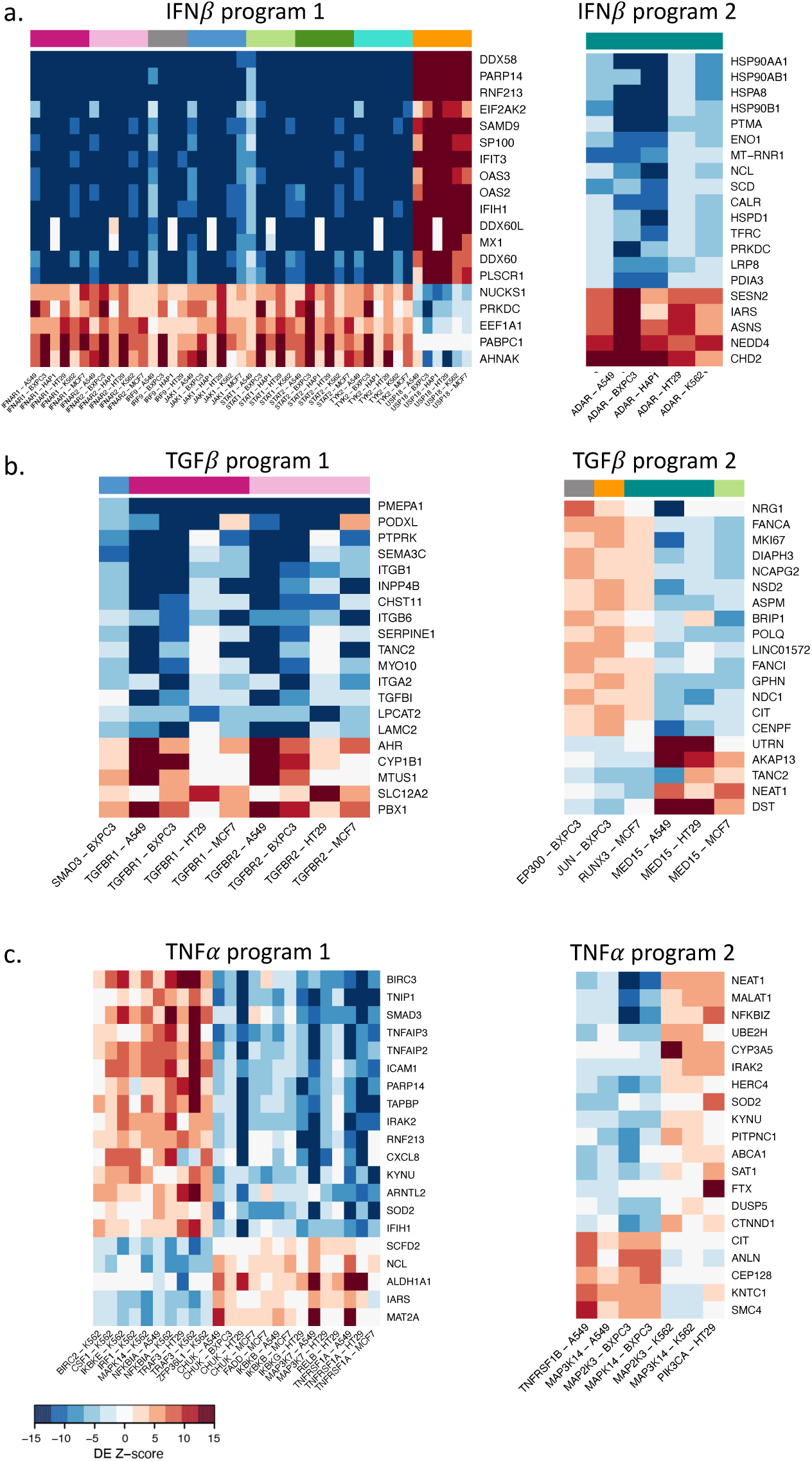
The first and second perturbation programs for IFNβ, TGFβ, and TNFα pathways, identified by MultiCCA. Each panel **(a)** IFNβ, **(b)** TGFβ, and **(c)** TNFα, shows a heatmap where columns represent correlated perturbations within and across cell lines, and rows list the program’s top 15 down-regulated genes and top 5 up-regulated genes. As in Figure 3g, the color gradient in the heatmap cells reflects the DE test Z-scores for each gene under each perturbation. See Supplementary Table 3 for a complete lists of pathway programs and the corresponding program genes.

**Supplementary Figure 7.**
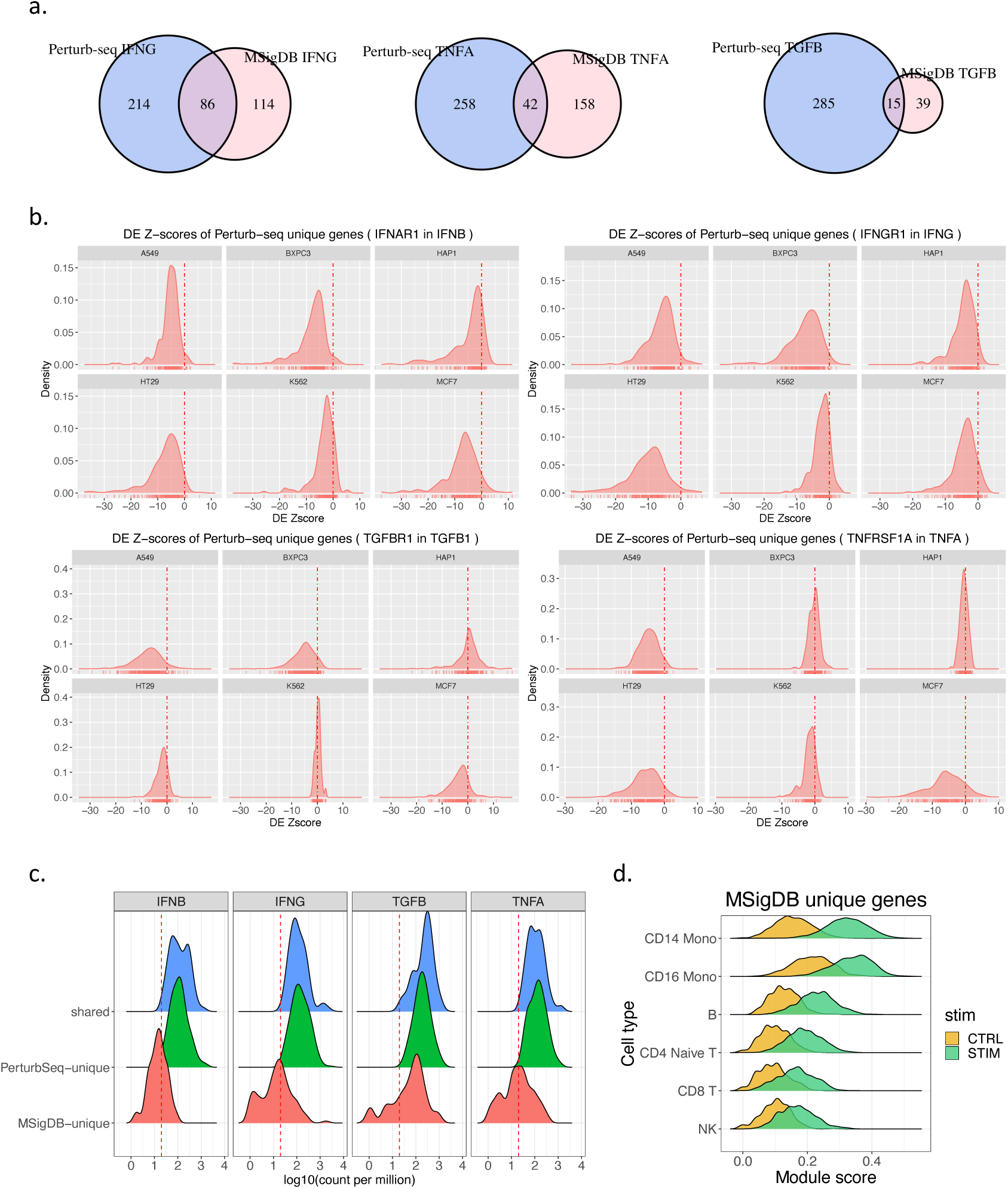
**(a).** Venn diagrams showing the overlap between the MultiCCA program 1 genes we identified and the MSigDB Hallmark gene lists for IFNγ, TGFβ, and TNFα pathways (Supplementary Methods). **(b).** Density plot for the distributions of DE Z-scores for the Perturb-seq unique genes. The density plots are generated using the perturbations from a key regulator in each pathway (labeled in the figure titles) using our Perturb-seq datasets. Plot shows that the uniquely identified Perturb-seq genes are repeatedly identified across multiple cell lines **(c).** Density plot for the log_10_(count per million) of MSigDB-unique, Perturb-seq-unique, and shared genes for each pathway, calculated in our Perturb-seq data (pseudobulk for all cells). The red dashed line indicates log_10_(CPM=20). Plot shows that genes identified by MSigDB have a very different expression profile than those either unique identified by Perturb-seq or shared between the two databases **(d).** IFNβ module score comparing unstimulated and stimulated cells using the MSigDB unique gene set. Plot shows that the MSigDB-unique gene sets effectively discriminates stimulated and control cells in only some cell types, in contrast with Perturb-seq gene sets in Figure 4c.

**Supplementary Figure 8.**
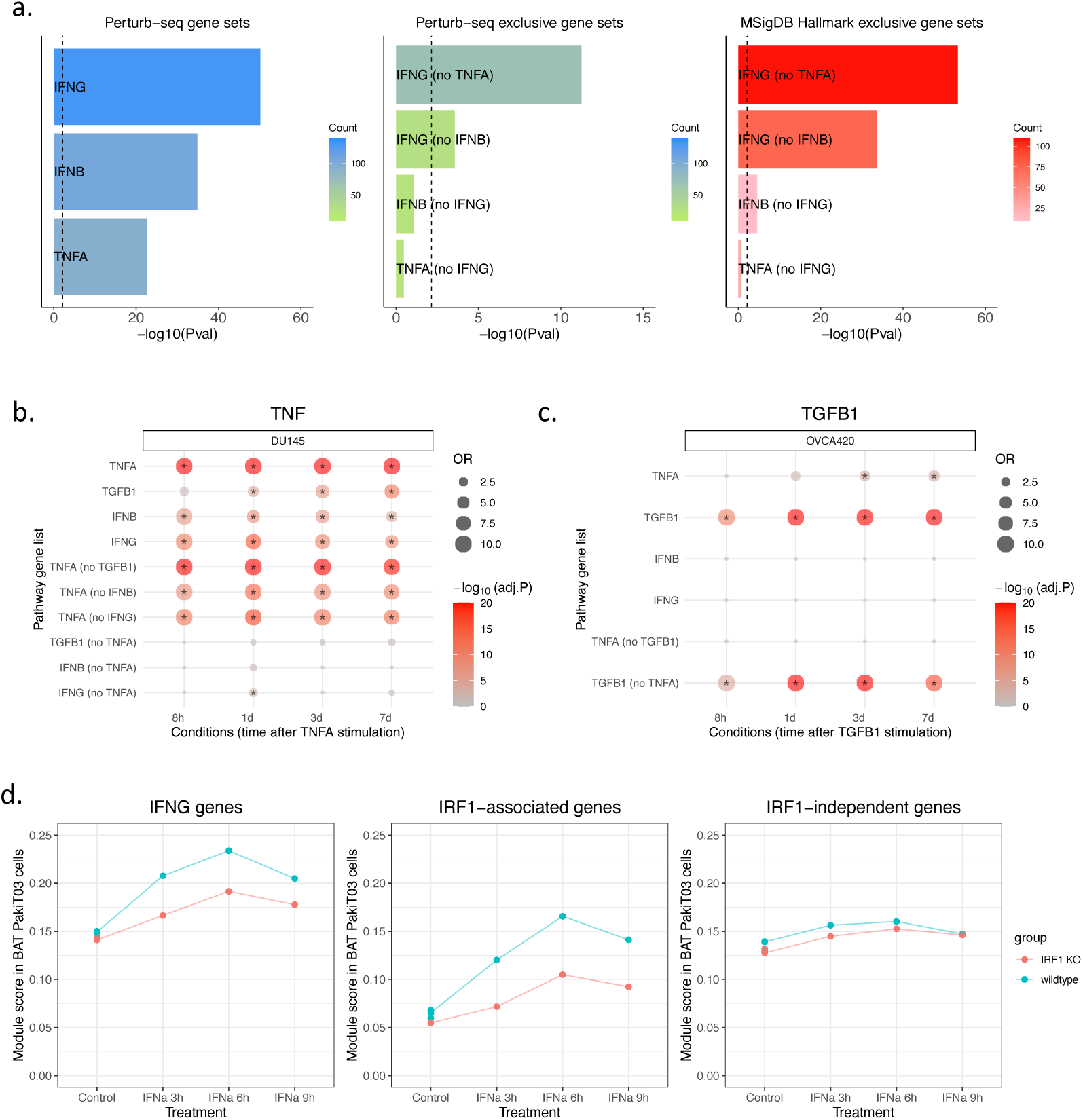
**(a-c).** Evaluating complete pathway and pathway-exclusive gene sets. Plots are as in Figure 4b, but run on datasets of cells that are stimulated with a single cytokine as a positive control. (a) shows the results from human CD14 monocytes stimulated with IFNγ, (b) shows results from the DU145 cell line stimulated with TNFα, and (c) shows results from the OVCA420 cells stimulated with TGFβ (Supplementary Methods). In each case, our Perturb-seq pathway lists show enrichment, but there is also enriched signal for alternative pathways since pathway gene sets include overlapping genes. Once we restrict the analysis to pathway exclusive gene sets, only the correct pathway exhibits evidence of enrichment. **(d).** The module score for IFNγ pathway genes, IRF1-associated genes, and IRF1-independent genes calculated in an IRF1-KO bat PakiT03 cell dataset (Supplementary Methods). The IRF1-associated genes and IRF1-independent genes are identified using the IFNγ program 1 and 2 in our Perturb-seq data (Supplementary Methods).

**Supplementary Figure 9.**
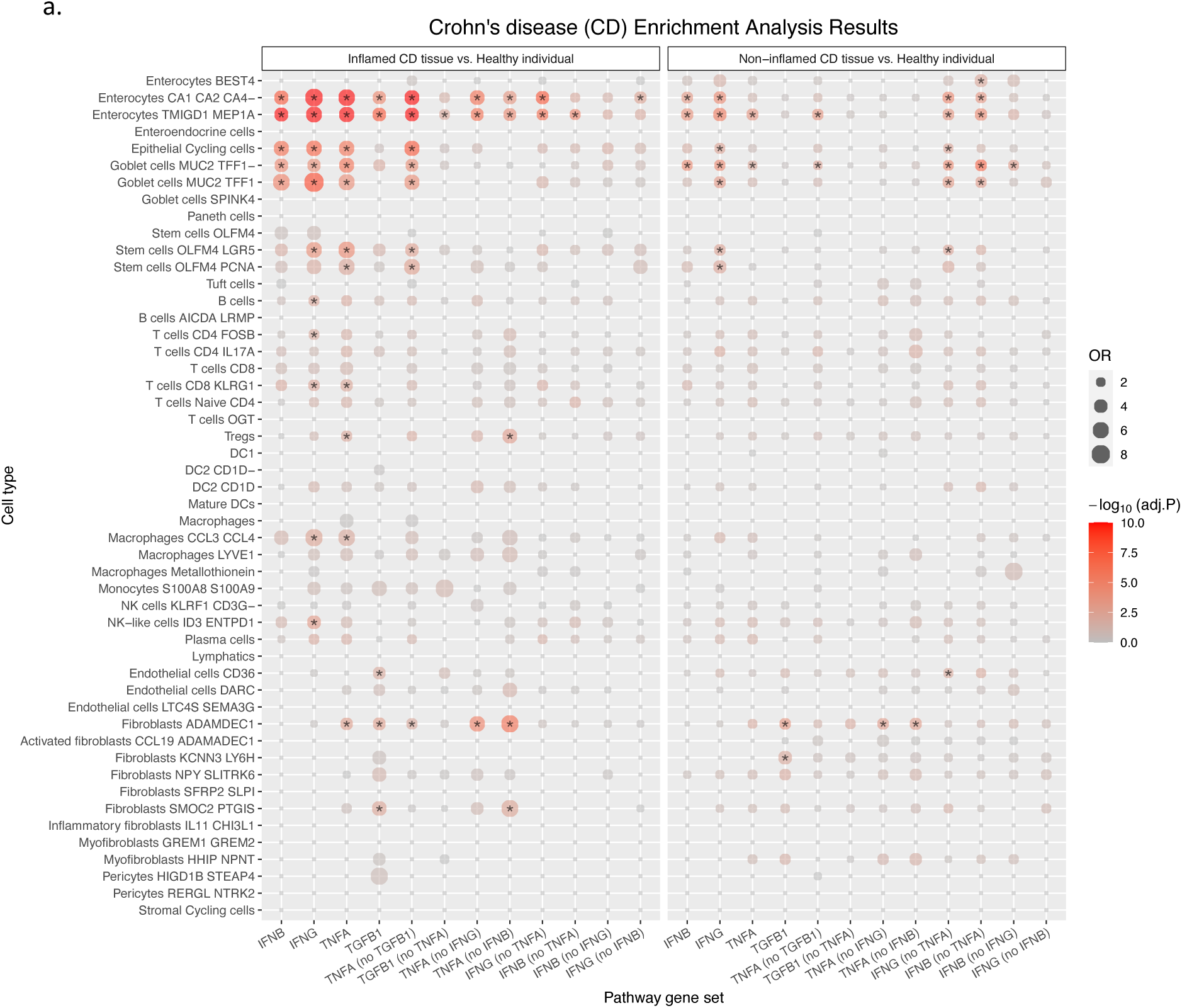
**(a).** The gene set enrichment test for DEG identified for patients with Crohn’s disease (CD) in an external dataset (Supplementary Methods). The analysis includes inflamed and non-inflamed tissues from CD patients. Each row indicates a gene set from our Perturb-seq data, and each column indicates a cell type from which the DEGs are obtained. The enrichment test odds ratio is represented by the size of the dot, and the enrichment test adjusted P-value (after Benjamini-Hochberg correction) is represented by the gradient of the color. Adjusted P-values less than 0.01 are labelled by asterisk.

## REFERENCES

1. Tang, F. et al. mRNA-Seq whole-transcriptome analysis of a single cell. Nat. Methods 6, 377–382 (2009).

2. Macosko, E. Z. et al. Highly Parallel Genome-wide Expression Profiling of Individual Cells Using Nanoliter Droplets. Cell 161, 1202–1214 (2015).

3. Picelli, S. et al. Smart-seq2 for sensitive full-length transcriptome profiling in single cells. Nat. Methods 10, 1096–1098 (2013).

4. Buenrostro, J. D., Wu, B., Chang, H. Y. & Greenleaf, W. J. ATAC-seq: A Method for Assaying Chromatin Accessibility Genome-Wide. Curr. Protoc. Mol. Biol. 109, 21.29.1–21.29.9 (2015).

5. Farlik, M. et al. Single-cell DNA methylome sequencing and bioinformatic inference of epigenomic cell-state dynamics. Cell Rep. 10, 1386–1397 (2015).

6. Stoeckius, M. et al. Simultaneous epitope and transcriptome measurement in single cells. Nat. Methods 14, 865–868 (2017).

7. Rotem, A. et al. Single-cell ChIP-seq reveals cell subpopulations defined by chromatin state. Nat. Biotechnol. 33, 1165–1172 (2015).

8. Mimitou, E. P. et al. Scalable, multimodal profiling of chromatin accessibility, gene expression and protein levels in single cells. Nat. Biotechnol. 39, 1246–1258 (2021).

9. Stubbington, M. J. T., Rozenblatt-Rosen, O., Regev, A. & Teichmann, S. A. Single-cell transcriptomics to explore the immune system in health and disease. Science 358, 58–63 (2017).

10. Van Hove, H. et al. A single-cell atlas of mouse brain macrophages reveals unique transcriptional identities shaped by ontogeny and tissue environment. Nat. Neurosci. 22, 1021–1035 (2019).

11. Velmeshev, D. et al. Single-cell genomics identifies cell type-specific molecular changes in autism. Science 364, 685–689 (2019).

12. Srivatsan, S. R. et al. Massively multiplex chemical transcriptomics at single-cell resolution. Science 367, 45–51 (2020).

13. Mulder, K. et al. Cross-tissue single-cell landscape of human monocytes and macrophages in health and disease. Immunity 54, 1883–1900.e5 (2021).

14. Tabula Sapiens Consortium* et al. The Tabula Sapiens: A multiple-organ, single-cell transcriptomic atlas of humans. *Science* 376, eabl4896 (2022).

15. Jinek, M. et al. A programmable dual-RNA-guided DNA endonuclease in adaptive bacterial immunity. Science 337, 816–821 (2012).

16. Cong, L. et al. Multiplex genome engineering using CRISPR/Cas systems. Science 339, 819–823 (2013).

17. Shalem, O. et al. Genome-scale CRISPR-Cas9 knockout screening in human cells. Science 343, 84– 87 (2014).

18. Wang, T., Wei, J. J., Sabatini, D. M. & Lander, E. S. Genetic screens in human cells using the CRISPR-Cas9 system. Science 343, 80–84 (2014).

19. Li, W. et al. MAGeCK enables robust identification of essential genes from genome-scale CRISPR/Cas9 knockout screens. Genome Biol. 15, 554 (2014).

20. Dixit, A. et al. Perturb-Seq: Dissecting Molecular Circuits with Scalable Single-Cell RNA Profiling of Pooled Genetic Screens. Cell 167, 1853–1866.e17 (2016).

21. Adamson, B. et al. A Multiplexed Single-Cell CRISPR Screening Platform Enables Systematic Dissection of the Unfolded Protein Response. Cell 167, 1867–1882.e21 (2016).

22. Jaitin, D. A. et al. Dissecting Immune Circuits by Linking CRISPR-Pooled Screens with Single-Cell RNA-Seq. Cell 167, 1883–1896.e15 (2016).

23. Shifrut, E. et al. Genome-wide CRISPR Screens in Primary Human T Cells Reveal Key Regulators of Immune Function. Cell 175, 1958–1971.e15 (2018).

24. Gasperini, M. et al. A Genome-wide Framework for Mapping Gene Regulation via Cellular Genetic Screens. Cell 176, 377–390.e19 (2019).

25. Replogle, J. M. et al. Mapping information-rich genotype-phenotype landscapes with genome-scale Perturb-seq. Cell 185, 2559–2575.e28 (2022).

26. The Gene Ontology Consortium. Expansion of the Gene Ontology knowledgebase and resources. *Nucleic Acids Res.* 45, D331–D338 (2017).

27. Jassal, B. et al. The reactome pathway knowledgebase. Nucleic Acids Res. 48, D498–D503 (2020).

28. Kanehisa, M., Furumichi, M., Tanabe, M., Sato, Y. & Morishima, K. KEGG: new perspectives on genomes, pathways, diseases and drugs. Nucleic Acids Res. 45, D353–D361 (2017).

29. Liberzon, A. et al. The Molecular Signatures Database (MSigDB) hallmark gene set collection. Cell Syst 1, 417–425 (2015).

30. Morris, J. A. et al. Discovery of target genes and pathways at GWAS loci by pooled single-cell CRISPR screens. Science 380, eadh7699 (2023).

31. Yeo, N. C. et al. An enhanced CRISPR repressor for targeted mammalian gene regulation. Nat. Methods 15, 611–616 (2018).

32. Börold, J. et al. BRD9 is a druggable component of interferon-stimulated gene expression and antiviral activity. EMBO Rep. 22, e52823 (2021).

33. Banno, T., Gazel, A. & Blumenberg, M. Effects of Tumor Necrosis Factor-α (TNFα) in Epidermal Keratinocytes Revealed Using Global Transcriptional Profiling*. J. Biol. Chem. 279, 32633–32642 (2004).

34. Zeng, C.-M., Chen, Z. & Fu, L. Frizzled Receptors as Potential Therapeutic Targets in Human Cancers. Int. J. Mol. Sci. 19, (2018).

35. Sato, M. et al. Distinct and essential roles of transcription factors IRF-3 and IRF-7 in response to viruses for IFN-alpha/beta gene induction. Immunity 13, 539–548 (2000).

36. Kubiczkova, L., Sedlarikova, L., Hajek, R. & Sevcikova, S. TGF-β - an excellent servant but a bad master. J. Transl. Med. 10, 183 (2012).

37. Barkett, M. & Gilmore, T. D. Control of apoptosis by Rel/NF-kappaB transcription factors. Oncogene 18, 6910–6924 (1999).

38. Honda, K., Takaoka, A. & Taniguchi, T. Type I interferon [corrected] gene induction by the interferon regulatory factor family of transcription factors. Immunity 25, 349–360 (2006).

39. Gordon, M. D. & Nusse, R. Wnt Signaling: Multiple Pathways, Multiple Receptors, and Multiple Transcription Factors*. J. Biol. Chem. 281, 22429–22433 (2006).

40. Pico, A. R. et al. WikiPathways: pathway editing for the people. PLoS Biol. 6, e184 (2008).

41. Sanson, K. R. et al. Optimized libraries for CRISPR-Cas9 genetic screens with multiple modalities. Nat. Commun. 9, 5416 (2018).

42. Tran, V. et al. High sensitivity single cell RNA sequencing with split pool barcoding. *bioRxiv* 2022.08.27.505512 (2022) doi:10.1101/2022.08.27.505512.

43. Almogy, G. et al. Cost-efficient whole genome-sequencing using novel mostly natural sequencing-by-synthesis chemistry and open fluidics platform. *bioRxiv* 2022.05.29.493900 (2022) doi:10.1101/2022.05.29.493900.

44. Squair, J. W. et al. Confronting false discoveries in single-cell differential expression. Nat. Commun. 12, 5692 (2021).

45. Wang, T., Li, B., Nelson, C. E. & Nabavi, S. Comparative analysis of differential gene expression analysis tools for single-cell RNA sequencing data. BMC Bioinformatics 20, 40 (2019).

46. Papalexi, E. et al. Characterizing the molecular regulation of inhibitory immune checkpoints with multimodal single-cell screens. Nat. Genet. 53, 322–331 (2021).

47. Replogle, J. M. et al. Maximizing CRISPRi efficacy and accessibility with dual-sgRNA libraries and optimal effectors. Elife 11, (2022).

48. Alerasool, N., Segal, D., Lee, H. & Taipale, M. An efficient KRAB domain for CRISPRi applications in human cells. Nat. Methods 17, 1093–1096 (2020).

49. Gilbert, L. A. et al. Genome-Scale CRISPR-Mediated Control of Gene Repression and Activation. Cell 159, 647–661 (2014).

50. Xu, H. et al. Sequence determinants of improved CRISPR sgRNA design. Genome Res. 25, 1147– 1157 (2015).

51. Jost, M. et al. Titrating gene expression using libraries of systematically attenuated CRISPR guide RNAs. Nat. Biotechnol. 38, 355–364 (2020).

52. Simmons, S. K. et al. Mostly natural sequencing-by-synthesis for scRNA-seq using Ultima sequencing. Nat. Biotechnol. 41, 204–211 (2023).

53. Ivashkiv, L. B. IFNγ: signalling, epigenetics and roles in immunity, metabolism, disease and cancer immunotherapy. Nat. Rev. Immunol. 18, 545–558 (2018).

54. Witten, D. M., Tibshirani, R. & Hastie, T. A penalized matrix decomposition, with applications to sparse principal components and canonical correlation analysis. Biostatistics 10, 515–534 (2009).

55. Jerby-Arnon, L. & Regev, A. DIALOGUE maps multicellular programs in tissue from single-cell or spatial transcriptomics data. Nat. Biotechnol. 40, 1467–1477 (2022).

56. François-Newton, V. et al. USP18-based negative feedback control is induced by type I and type III interferons and specifically inactivates interferon α response. PLoS One 6, e22200 (2011).

57. Basters, A., Knobeloch, K.-P. & Fritz, G. USP18 - a multifunctional component in the interferon response. Biosci. Rep. 38, (2018).

58. Oshima, S. et al. Interferon regulatory factor 1 (IRF-1) and IRF-2 distinctively up-regulate gene expression and production of interleukin-7 in human intestinal epithelial cells. Mol. Cell. Biol. 24, 6298–6310 (2004).

59. Harada, H. et al. Structurally similar but functionally distinct factors, IRF-1 and IRF-2, bind to the same regulatory elements of IFN and IFN-inducible genes. Cell 58, 729–739 (1989).

60. Bien, J. & Tibshirani, R. Hierarchical Clustering With Prototypes via Minimax Linkage. J. Am. Stat. Assoc. 106, 1075–1084 (2011).

61. Alsamman, K. & El-Masry, O. S. Interferon regulatory factor 1 inactivation in human cancer. Biosci. Rep. 38, (2018).

62. Pollaci, G. et al. Novel Multifaceted Roles for RNF213 Protein. Int. J. Mol. Sci. 23, (2022).

63. Grünvogel, O. et al. DDX60L Is an Interferon-Stimulated Gene Product Restricting Hepatitis C Virus Replication in Cell Culture. J. Virol. 89, 10548–10568 (2015).

64. Kang, H. M. et al. Multiplexed droplet single-cell RNA-sequencing using natural genetic variation. Nat. Biotechnol. 36, 89–94 (2018).

65. Kartha, V. K. et al. Functional inference of gene regulation using single-cell multi-omics. Cell Genom 2, (2022).

66. Cook, D. P. & Vanderhyden, B. C. Context specificity of the EMT transcriptional response. Nat. Commun. 11, 2142 (2020).

67. Rosain, J. et al. Human IRF1 governs macrophagic IFN-γ immunity to mycobacteria. Cell 186, 621– 645.e33 (2023).

68. Irving, A. T. et al. Interferon Regulatory Factors IRF1 and IRF7 Directly Regulate Gene Expression in Bats in Response to Viral Infection. Cell Rep. 33, 108345 (2020).

69. Lei, X. et al. Activation and evasion of type I interferon responses by SARS-CoV-2. Nat. Commun. 11, 3810 (2020).

70. Hadjadj, J. et al. Impaired type I interferon activity and inflammatory responses in severe COVID-19 patients. Science 369, 718–724 (2020).

71. Lee, J. S., et al. Immunophenotyping of COVID-19 and influenza highlights the role of type I interferons in development of severe COVID-19. *Sci Immunol* 5, (2020).

72. Toro, A. et al. Pin-Pointing the Key Hubs in the IFN-γ Pathway Responding to SARS-CoV-2 Infection. Viruses 14, (2022).

73. Gadotti, A. C. et al. IFN-γ is an independent risk factor associated with mortality in patients with moderate and severe COVID-19 infection. Virus Res. 289, 198171 (2020).

74. Karki, R. et al. Synergism of TNF-α and IFN-γ Triggers Inflammatory Cell Death, Tissue Damage, and Mortality in SARS-CoV-2 Infection and Cytokine Shock Syndromes. *Cell* **184**, 149–168.e17 (2021).

75. COvid-19 Multi-omics Blood ATlas (COMBAT) Consortium. Electronic address: & COvid-19 Multi-omics Blood ATlas (COMBAT) Consortium. A blood atlas of COVID-19 defines hallmarks of disease severity and specificity. Cell 185, 916– 938.e58 (2022).

76. Kong, L. et al. The landscape of immune dysregulation in Crohn’s disease revealed through single-cell transcriptomic profiling in the ileum and colon. Immunity 56, 444–458.e5 (2023).

77. Liu, T.-C. & Stappenbeck, T. S. Genetics and Pathogenesis of Inflammatory Bowel Disease. Annu. Rev. Pathol. 11, 127–148 (2016).

78. Parigi, S. M. et al. The spatial transcriptomic landscape of the healing mouse intestine following damage. Nat. Commun. 13, 828 (2022).

79. Schubert, M. et al. Perturbation-response genes reveal signaling footprints in cancer gene expression. Nat. Commun. 9, 20 (2018).

80. Beck, P. L. et al. Transforming growth factor-beta mediates intestinal healing and susceptibility to injury in vitro and in vivo through epithelial cells. Am. J. Pathol. 162, 597–608 (2003).

81. Oshima, H. et al. Suppressing TGFβ signaling in regenerating epithelia in an inflammatory microenvironment is sufficient to cause invasive intestinal cancer. Cancer Res. 75, 766–776 (2015).

82. Penn, J. W., Grobbelaar, A. O. & Rolfe, K. J. The role of the TGF-β family in wound healing, burns and scarring: a review. Int. J. Burns Trauma 2, 18–28 (2012).

83. Datlinger, P. et al. Ultra-high-throughput single-cell RNA sequencing and perturbation screening with combinatorial fluidic indexing. Nat. Methods 18, 635–642 (2021).

84. Xu, Z., Sziraki, A., Lee, J., Zhou, W. & Cao, J. Dissecting key regulators of transcriptome kinetics through scalable single-cell RNA profiling of pooled CRISPR screens. Nat. Biotechnol. (2023) doi:10.1038/s41587-023-01948-9.

85. Lamb, J. et al. The Connectivity Map: using gene-expression signatures to connect small molecules, genes, and disease. Science 313, 1929–1935 (2006).

86. Calderon, D. et al. Landscape of stimulation-responsive chromatin across diverse human immune cells. Nat. Genet. 51, 1494–1505 (2019).

87. Urrutia, A. et al. Standardized Whole-Blood Transcriptional Profiling Enables the Deconvolution of Complex Induced Immune Responses. Cell Rep. 16, 2777–2791 (2016).

88. Jiang, P. et al. Systematic investigation of cytokine signaling activity at the tissue and single-cell levels. Nat. Methods 18, 1181–1191 (2021).

89. Cui, A. et al. Dictionary of immune responses to cytokines at single-cell resolution. Nature (2023) doi:10.1038/s41586-023-06816-9.

90. Xie, Y. et al. Comparative analysis of single-cell RNA sequencing methods with and without sample multiplexing. bioRxiv 2023.06.28.546827 (2023) doi:10.1101/2023.06.28.546827.

91. Rubin, A. J. et al. Coupled Single-Cell CRISPR Screening and Epigenomic Profiling Reveals Causal Gene Regulatory Networks. Cell 176, 361–376.e17 (2019).

92. Liscovitch-Brauer, N. et al. Profiling the genetic determinants of chromatin accessibility with scalable single-cell CRISPR screens. Nat. Biotechnol. 39, 1270–1277 (2021).

93. Pierce, S. E., Granja, J. M. & Greenleaf, W. J. High-throughput single-cell chromatin accessibility CRISPR screens enable unbiased identification of regulatory networks in cancer. Nat. Commun. 12, 2969 (2021).

94. Mimitou, E. P. et al. Multiplexed detection of proteins, transcriptomes, clonotypes and CRISPR perturbations in single cells. Nat. Methods 16, 409–412 (2019).

95. Wessels, H.-H. et al. Efficient combinatorial targeting of RNA transcripts in single cells with Cas13 RNA Perturb-seq. Nat. Methods 20, 86–94 (2023).

96. Jiang, L. et al. Systematic reconstruction of molecular pathway signatures using massively scalable single-cell perturbation screens. (2024) doi:10.5281/ZENODO.10520190.

