## Supplementary Methods for "Systematic reconstruction of molecular pathway signatures using scalable single-cell perturbation screens"

#### Cell culture and monoclonal CRISPRi cell line generation

HEK293FT cells were acquired from Thermo Fisher (R70007), A549 cells were acquired from ATCC (CCL-185), BXPC3 cells were acquired from ATCC (CRL-1687), HAP1 cells were acquired from Horizon Discovery (C631), HT29 cells were acquired from ATCC (HTB-38), K562 cells were acquired from ATCC (CCL-243), and MCF7 cells were acquired from ATCC (HTB-22). HEK293FT and A549 cells were grown in DMEM with high glucose with L-glutamine and sodium pyruvate (Cytiva SH30243.01), BXPC3 cells were grown in RPMI 1640 (Caisson Labs RPL03), HAP1 and K562 cells were grown in IMDM with GlutaMAX™ Supplement (Thermo Scientific 31980030), HT29 cells were grown in McCoy's 5A (Caisson Labs MCL02), and MCF7 cells were grown in EMEM (Sigma-Aldrich M4655). All media was supplemented with 10% fetal bovine serum (Corning 35-010-CV) and all cell lines were maintained at 37°C with 5% $CO_2$.

Monoclonal dCas9-KRAB-MeCP2 A549, BxPC3, HAP1, HT29, and MCF7 cell lines were prepared as previously described [1], and the K562 monoclonal dCas9-KRAB-MeCP2 cell line was a gift from the Sanjana Lab. Briefly, cells were lentivirally transduced with lentiCRISPRi(v2)-Blast (Addgene #170068) and then diluted and replated to isolate individual clones. The dCas9-KRAB-MeCP2 effector efficiency in each monoclonal line was evaluated first by performing a Western Blot for the Cas9 protein and second by transducing cells with a single guide RNA targeting the CD55 locus, verifying knockdown by flow cytometry (data not shown).

#### CRISPR guide library cloning

Five separate guide pools, one per signaling pathway, were designed using three guides per gene from the Dolcetto CRISPRi guide library [2]. Signaling pathway specific forward and reverse PCR handles were added to each guide sequence so that all five guide pools could be ordered as one large oligonucleotide pool from Twist Biosciences (Supplementary Table 1) with the following format:

Pathway-PCR-handle::partial-hU6-promoter::gRNA::partial-guidescaffold

The oligonucleotide pool was amplified for cloning using two PCRs. The first PCR was a targeted amplification of guides for a single pathway using pathway-specific PCR primers that match the PCR handles of the oligos ordered in pool format. For each signaling pathway, 1 ng of pooled oligos was amplified with pathway-specific primers and Q5 polymerase (NEB M0491L) using the following PCR program: 98 °C for 30 s; 9 cycles of 98 °C for 10 s, 63°C for 10 s and 72°C for 15 s; followed by 72°C for 3 min. The second PCR added sequences that are homologous to the cloning vector to each end of the guide oligonucleotides. For each pathway, purified product from the previous PCR was amplified with cloning homology primers and Q5 polymerase (NEB M0491L) using the following PCR program: 98 °C for 30 s; 8 cycles of 98 °C for 10 s, 72°C for 10 s and 72°C for 15 s; followed by 72°C for 3 min. The PCR product was purified using a 2.5X SPRI cleanup. The amplified oligo pool was ligated into a digested CROP-seq cloning vector using a Gibson Assembly reaction (NEB E2611S), following manufacturer recommendations.

**Lentivirus transduction and stimulation**

For virus production, 10 million HEK293FT cells were seeded in a 10 cm dish for 12-18 hours before transfection. Per transfection reaction we used 45µl of 1 mg/mL polyethyleneimine (PEI, Polysciences 23966-1) and 20 µg of total plasmid DNA (6.4 µg psPAX2: Addgene #12260; 4.4 µg pMD2.G: Addgene #12259; 9.2 µg of sgRNA expressing plasmid). Six to eight hours post-transfection, the medium was exchanged for 10 mL of DMEM + 10% FBS containing 1% bovine serum albumin (VWR 97061-420). Viral supernatants were collected after an additional 48 hours, spun down to remove cellular debris for 5 min at 4°C and 1000xg, and stored at -80°C until use.

Samples were prepared one pathway at a time. All six CRISPRi cell lines were transduced at a low MOI (0.05 – 0.1) with lentivirus for a pathway-specific guide pool. After three to five days of selection for guide positive cells with 1 µg/mL puromycin (Thermo Scientific A1113803) and seven to ten days of expansion, cells were stimulated for 24 hours with the corresponding cytokine to induce signaling: IFNβ (PBL Assay Science 11415-1), IFNγ (R&D Systems 285-IF-100), TGFβ (Thermo Scientific PHG9204), TNFα (R&D Systems 210-TA-005), or insulin (INS) (Sigma-Aldrich 91077C-250MG). Samples were then fixed using the Parse Biosciences Cell Fixation kit (ECF100) and stored at -80°C.

In total, our study comprised 30 biological contexts (six cell lines × five signaling pathways). After fixation, we split cells into two, creating a total of 60 samples that encompass two scRNA-seq replicates.

**Parse scRNA-seq with guide capture**

Fixed samples were thawed and single-cell RNA-seq with CRISPR guide capture was performed using the Parse Biosciences Evercode™ Whole Transcriptome v1 (ECW01030) or Whole Transcriptome Mega v1 (ECW01050) kit with minor modifications to the Parse protocol. Briefly, polyadenylated gRNAs were reverse transcribed and barcoded using the existing primers in the commercial kit, enriched during cDNA amplification using a guide additive primer, and amplified for sequencing using a two-step guide-specific library preparation protocol

(Supplementary Figure 1, Supplementary File 1). We optimized the guide capture conditions through guide additive cDNA amplification primer design (Supplementary Figure 2a-c), modifying the PCR annealing temperature to perform all cycles at 65°C (Supplementary Figure 2d-e), optimizing the guide additive primer concentration (Supplementary Figure 2f-g), and increasing the SPRI cleanup ratio from 0.8X to 1X to preserve shorter random hexamer reverse transcription derived guides (Supplementary Figure 2h).

### Next-generation sequencing

Our study comprises 60 different samples (6 cell lines × 5 signaling pathways × 2 replicates). gRNA libraries for all samples were sequenced using the Illumina NovaSeq platform. For gene expression libraries, we sequenced all libraries from the first replicate with Illumina NovaSeq. For a subset of 12 samples (IFNβ, INS pathway cells), we also sequenced samples using the Ultima UG100 platform. To perform Ultima UG100 sequencing, Illumina-compatible libraries were shipped to Ultima Genomics for library conversion following a previously described procedure [3], where Illumina adaptors are exchanged for Ultima-compatible adaptors using a PCR-based library conversion process. When sequencing scRNA-seq libraries from Parse Biosciences with Illumina, we use paired-end sequencing, with one read capturing cellular and molecular barcode information (barcode read), and the other read capturing the genetic sequence of the target molecule (cDNA read). When using Ultima sequencing, both barcode sequences and cDNA sequences are obtained in a single long read (200-250bp), which are separated by a poly(dT) stretch, and sequenced as previously described [3]. After we confirmed that Illumina sequencing and Ultima sequencing returned highly concordant differentially expressed gene sets (Supplementary Figure 5), we proceeded to sequence all of the second replicate samples with the Ultima UG100 platform.

### Perturb-seq data pre-processing and gRNA calling

We processed Perturb-seq gene expression sequence data using the Parse split-pipe pipeline (version 0.9.6 available from https://support.parsebiosciences.com/hc/en-us). The pipeline performs barcode correction, read alignment, transcriptomic quantification, and cell filtration. Sequencing reads from the mRNA library were aligned to the human reference genome GRCh38.p12 (Ensembl release 93). Sequencing reads from the gRNA library were aligned to a reference consisting of both the human reference genome and a reference of gRNA sequences that we used in this experiment (Supplementary Table 1).

The Parse split-pipe pipeline takes paired-end read data (generated by the Illumina platform) as input, so we implemented a data conversion step to generate synthetic paired-end reads from the Ultima data [3]. The barcode read was created by extracting the barcode sequences and the linker sequences from the UG100 read. The cDNA read represented the sequence that followed the poly(dT) stretch in the UG100 read. We discarded the poly(dT) stretch, as well as the 5 ensuing bases. We took the reverse complement of the remaining sequence, performed quality trimming as in [3], and trimmed the final read to 120bp. The output of the Parse split-pipe pipeline is a gene expression matrix, filtered to remove low-quality cells. We loaded these matrices in Seurat v4 [4], and removed cells with low number of genes detected (< 500) or high mitochondrial gene content (> 20%)

We applied the following procedure to assign gRNA identity to individual cells. We first removed any cell where no gRNA reads were detected, any cell where more than five separate gRNA were detected, and any cell where the most abundant gRNA counts were less than 1.5 times the counts of the second-most abundant gRNA. We then initially assigned cells to the most abundantly detected gRNA. To refine these calls, we aimed to establish a threshold for each gRNA that would distinguish between true signal and ambient RNA contamination or barcode misassignment, and did this individually for each gRNA. We leveraged the fact that for each gRNA (for example, gRNA targeting the gene IFNGR1), our multiplexed dataset contained cells that were targeted with libraries that did not include this gene (i.e. TNF-stimulated cells were not exposed to IFNGR1 gRNA). For each gRNA, we considered the distribution of these 'negative' cells, and selected the 80th quantile of this distribution as a threshold. Cells where the most abundantly expressed gRNA did not clear this threshold were reassigned as Negative. This process, repeated for each gRNA, resulted in a binary "gRNA identity matrix" (0s and 1s). We retained all cells that were assigned one gRNA identity. For the major analyses in this manuscript, we pooled cells from both replicates together, and we assessed replicate reproducibility in Figure 2e and Supplementary Figure 4d-f.

**Removing confounding variation in Perturb-seq experiment**

After quality control and gRNA assignment, we performed standard log-normalization (NormalizeData), variable gene identification (FindVariableFeatures) and principal component analysis (RunPCA) on the quantified gene expression data (RNA assay) using Seurat. To address confounding factors such as cell cycle stage and cellular stress responses, we employed Seurat's CalcPerturbSig() function developed in our previous study [5]. This process involved, for each cell, identifying the top 20 nearest neighbors from the non-targeting control cells using the top 40 principal components. These neighbors represent cells in a similar biological state to the target cell. We then subtracted the average expression of the 20 cells from the original expression profile of the target cell. This is performed for each cell, and the resulting vector isolates the transcriptomic impact of the cell's genetic perturbation, minimizing known and unknown confounders [5]. These vectors are stored in the PRTB assay in the Seurat object.

### Overview of the Mixscale scoring method

We developed Mixscale (https://github.com/longmanz/Mixscale), an extension of the original Mixscape method [5], to measure perturbation heterogeneity in individual cells. This method (RunMixscale() function) involves the following steps for each targeted regulator *r*. Note that in multi-cell line experiments, step 2-4 is carried out for each cell line separately to capture heterogeneous perturbation responses across biological contexts.

1) We conduct differential expression (DE) analysis between cells with gRNAs targeting regulator *r* and those with non-targeting (NT) gRNAs, using the RNA assay in Seurat. In multi-cell line experiments, NT cells are randomly sub-sampled to match the cell line proportion in the target group. We identify DE genes based on a Bonferroni-adjusted p-value below 0.05 and an absolute $\log_2$-fold-change over 0.2. The top 100 genes are selected if there are more than 100. On the other hand, if fewer than five genes are identified, we stop the process and assign a score of 1 (i.e., $\tilde{s}_i^r = 1$ in step 4) to all target group cells.

2) We define perturbation vectors $= [\boldsymbol{p_1^r}, \ldots, \boldsymbol{p_N^r}]$ for *N* cells with gRNA targeting regulator *r* and $= [\boldsymbol{p_1^{NT}}, \ldots, \boldsymbol{p_M^{NT}}]$) for *M* cells with NT gRNA. Each cell's perturbation vector length equals the number of identified DE genes in step 1, with expression values from the PRTB assay.

3) We calculate the average difference in perturbation vectors between targeted and NT cells ($\boldsymbol{\overline{p}^r} - \boldsymbol{\overline{p}^{NT}}$), where $\boldsymbol{\overline{p}^r} = \sum_{i=1}^{N} \boldsymbol{p_i^r} / N$ and $\boldsymbol{\overline{p}^{NT}} = \sum_{j=1}^{M} \boldsymbol{p_j^{NT}} / M$ are the average vectors for each group. We then project each cell's perturbation vector $\boldsymbol{p_i}$ onto this difference vector to obtain the perturbation score $s_i$ for cell *i* (either from the target or the NT group):

$$s_i^r = \boldsymbol{p_i^r} \times (\boldsymbol{\overline{p}^r} - \boldsymbol{\overline{p}^{NT}})$$

$$s_i^{NT} = \boldsymbol{p_i^{NT}} \times (\boldsymbol{\overline{p}^g} - \boldsymbol{\overline{p}^{NT}})$$

The $s_i$ represents the magnitude of expression shift of cell *i* from the NT cells due to perturbation. Higher $s_i$ values indicate a stronger perturbation response.

4) We further calculate the mean and standard deviation of the scores in NT cells, and used them to standardize the scores in target cells:

$$\mu^{NT} = \frac{\sum_{i=1}^{M} s_i^{NT}}{M}$$

$$\sigma^{NT} = \sqrt{\frac{\sum_{i=1}^{M} (s_i^{NT} - \mu^{NT})^2}{M - 1}}$$

$$\tilde{s}_i^r = \frac{s_i^r - \mu^{NT}}{\sigma^{NT}}$$

The standardized scores can be used as input for weighted DE analysis (see below), enabling cells with a stronger perturbation response to have a greater contribution to the identification of differentially expressed genes (DEGs). While we identified that this weighting scheme increases statistical power, it also establishes a dependency between the calculation of Mixscale scores and the downstream identification of DEG. As a result, genes that are themselves involved in the calculation of Mixscale scores will therefore be more likely to be identified as DE. To address this, for each cell, we calculated 'leave one out' (LOO) standardized Mixscale scores, where we repeated steps 2-4 but after leaving out one individual gene. For each regulator, when performing DE test (see below) for genes that were involved in the construction of Mixscale scores, we used these LOO scores as input.

**Differential expression analysis**

We incorporated Mixscale scores as weights into negative binomial regression for differential expression (DE) analysis. We term this weighted multivariate regression framework as "wmvReg". We apply wmvReg to identify the response genes for each regulator in each pathway independently (i.e. to identify the downstream targets of STAT2 perturbation after IFNβ stimulation). Since each perturbation is performed in multiple cell lines, we combine data from all cell lines together when fitting the regression model. We also estimate cell line-specific regression coefficients to address biological heterogeneity. The wmvReg procedure (Run_wmvRegDE() function) is implemented in the Mixscale package (available at https://github.com/longmanz/Mixscale), and described below.

We define the following:

$\boldsymbol{y} \in \mathbb{R}^{N+M}$: Vector of scRNA-seq expression counts matrix for a response gene. Includes all $N$ cells (across $L$ cell lines) that are assigned targeting guides for a specific regulator, as well as all $M$ cells assigned non-targeting control guides.

$\mathbf{X} \in \mathbb{R}^{(N+M)\times L}$: Matrix of cell line indicator variables. $\boldsymbol{X_{i,j}}$ is 1 if cell $i$ originates from cell line $j$, and is 0 otherwise.

$\mathbf{S} \in \mathbb{R}^{(N+M)\times L}$: Matrix of mixscale scores. $\boldsymbol{S_{i,j}}$ is set to the mixscale score $\tilde{s}_i^r$ if cell $i$ received a targeting gRNA for regulator $r$ and originates from cell line $j$, and is 0 otherwise.

$\boldsymbol{c} \in \mathbb{R}^{N+M}$: Vector of the $log_{10}$-transformed total count for each cell, used to adjust for library size differences.

We then fit a regression model of the form:

$$log(E(\boldsymbol{y})) = \mathbf{X}\boldsymbol{\alpha} + \mathbf{S}\boldsymbol{\beta} + \boldsymbol{c}\gamma$$

We estimate three terms:

$\boldsymbol{\alpha} \in \mathbb{R}^{L}$ represents a vector of the the $log_{10}$ baseline expression of the response gene in each of $L$ cell lines.

$\boldsymbol{\beta} \in \mathbb{R}^{L}$ is the primary term we are interested in, and represents a vector of regression coefficients estimated for each of $L$ cell lines. Regression coefficients that deviate from 0 indicate that the perturbed regulator results in differential expression of the response gene.

$\gamma$ is a scalar that accounts for the relationship between gene expression levels and cellular library size.

We perform model fitting using the glmGamPoi R package (version 1.10.2) [6].

While employing scores in DE tests can increase the statistical power to detect DE genes (Figure 2e), we recognized an issue: the scores, being calculated from a subset of genes, create an artificial correlation between $y$ and $\tilde{s}_i^r$. To address this, we adopted a different strategy for the DE tests of genes used in score calculations. For these tests, we used their corresponding leave-one-out (LOO) scores, as mentioned in the previous section. This approach significantly reduced the inflation of test statistics when the number of mis-specified genes is relatively small, which is usually true in real practice (Supplementary Figure 3c).

In addition, we conducted DE analyses using conventional methods (Figure 2e). We used FindMarkers() in Seurat to perform Wilcoxon rank sum test [7] between NT cells and target cells in each cell line separately. We also conducted standard unweighted negative binomial regression, coding NT cells as 0 and target cells as 1, using the glmGamPoi R package [6].

### Evaluating false postive rate and statistical power for DE testing

To assess the calibration of our weighted DE method's test statistics, we scrambled the gRNA and NT cell labels, and repeated the DE test, enabling us to estimate false positive rates (FPR, proportion of genes with p-values ≤ alpha). We conducted the simulation in two ways. In the first simulation (Supplementary Figure 4a), we used Mixscale scores that were calculated prior to shuffling labels. We then performed unweighted and weighted DE analysis, respectively, between shuffled NT and target. We gathered DE p-values and calculated FPRs at alpha levels of 0.05, 0.01, and 0.005 for both methods for comparison.

In the second simulation (Supplementary Figure 4b), we recalculated Mixscale scores after gRNA label shuffling. In each of our simulations, we found that after label shuffling, we could not calculate Mixscale scores under our standard procedure, since we could not identify >=5 significant DEG between shuffled NT and target cells. For these simulations, we removed this threshold, and selected the top 10, 30, 50, or 100 DE genes with the smallest DE p-values for score calculation, even if they did not pass the significance threshold. Following the first simulation's approach, we performed unweighted and weighted DE analysis, respectively, between shuffled NT and target cells for all null genes, collected DE p-values, and calculated FPRs at the same alpha levels for both methods for comparison. We show these results in Supplementary Figure 4b, and note that they represent a substantial overstatement of FPR under null conditions, since when applied to real data, we would fail to calculate Mixscale scores and the DE test would degrade to a standard weighted test.

To evaluate the statistical power of our weighted DE method, we used the replication rate of DE genes between the two scRNA-seq replicates. For each DE method (weighted, unweighted, and Wilcoxon rank sum test), we first performed DE analysis between NT and target cells from the first replicate, for each perturbation target in each pathway. We identified significant DE genes with raw p-values $\leq 1.67\times10^{-6}$ (equivalent to 0.05 after Bonferroni correction for 30,000 genes transcriptome-wise). We then performed the same DE analysis on cells from the second replicate, and calculated the replication rate, defined as the proportion of these significant DE genes from replicate 1 that also had p-values $\leq 1.67\times10^{-6}$ in replicate 2.

**Joint analysis of two replicates**

Our first scRNA-seq replicate included cells that were sequenced by both the Illumina NovaSeq and Ultima UG100 platforms (see above). We compared the pseudobulk counts for each gene between the two datasets, identifying the top 20 genes with the most divergent counts as outliers (Supplementary Figure 5a). We then performed a weighted DE analysis separately on the Illumina and Ultima data, and identified high reproducibility of DEG p-values across sequencing technologies (Supplementary Figure 5b). Additionally, for each pathway, we extracted and plotted the distribution of DE z-scores for the 20 outlier genes across different perturbation groups and cell lines (Supplementary Figure 5c), confirming that while these genes exhibited differential gene counts between Illumina and Ultima data, they did not exhibit differential expression when comparing NT and targeted cells within a sequencing platform. Based on these results, we sequenced cells from our second scRNA-seq replicate with the Ultima UG100 platform, and pooled the two replicates together for joint analysis.

#### Decomposition analysis to identify correlated perturbations

We adapted the DIALOGUE method [8], a MultiCCA-based [9] decomposition approach, to identify shared DE gene patterns across perturbations and cell lines within each pathway (Figure 3e). The following steps were performed for each pathway individually (codes implemented in Mixscale R package, available at https://github.com/longmanz/Mixscale).

After completing the DE analysis, we perform the following decomposition analysis for each pathway independently. We define the following for each perturbed regulator *r*, each cell line *l*, and each response gene $g$:

$\beta_{r,l,g}$: regression coefficient from the DE test

$\sigma_{r,l,g}$: standard error of the regression coefficient

$z_{r,l,g} = \frac{\beta_{r,l,g}}{\sigma_{r,l,g}}$ : z-score for the regression coefficient

1) We organize the z-scores into a tensor $\boldsymbol{P} \in \mathbb{R}^{R \times L \times G}$, where $R$ is the total number of perturbed regulators, $L$ is the total number of cell lines, and $G$ represents the total number of potential response genes. The value of $G$ fluctuates between 8,169 and 8,954, depending on the pathway. Missing entries in $\boldsymbol{P}$ are filled with 0.

2) Next, we divide the tensor $\boldsymbol{P}$ along its second dimension into a set of matrices $\{\mathbf{M_1}, \dots, \mathbf{M}_L\}$, where $\mathbf{M_i} \in \mathbb{R}^{R \times G}$ represents the DE Z-scores of the *i-th* cell line. From these matrices, we selected gene set $G'$, representing the union of the top 300 significant DE genes for each individual regulator. The size of $G'$ ranges from 1,314 to 2,486 depending on the pathway, and helps to focus the downstream analysis on the most relevant set of DEG. We then define the filtered matrices as $\{\mathbf{Q_1}, \dots, \mathbf{Q}_L\}$ where $\mathbf{Q_i} \in \mathbb{R}^{R \times G'}$.

3) Given matrices $\{\mathbf{Q_1}, \dots, \mathbf{Q}_L\}$, we apply MultiCCA (implemented in the PMA R package [9], version 1.2.1) to search for a series of $R$×1 transformation vectors $w_1$, ..., $w_L$ (canonical weights) that solves the following optimization problem [9]:

$$maximize \sum_{i<j} w_i^T \mathbf{Q_i^T} \mathbf{Q_j} w_j \, , subject\ to\ \forall i\ \|w_i\| \leq 1, p_i(w_i) \leq c_i$$

Where $p_i(w_i)$ is the LASSO penalties. The equation can be interpreted as maximizing the sum of pairwise correlations of the canonical vectors ($cv_i = \mathbf{Q_i}w_i$ and $cv_j = \mathbf{Q_j}w_j$) across all $\{\mathbf{Q_1}, \dots, \mathbf{Q}_L\}$. In our analyses, we did not apply LASSO penalties to force $w_i$ to be sparse.

4) After determining the optimized $w_i$ for each $\mathbf{Q_i}$, we calculate the canonical vector $cv_i$ ($= \mathbf{Q_i}w_i$). This vector describes a downstream perturbation signal in the $i$-th cell shared across multiple regulators. We associated specific regulator/cell lines combinations (columns of $\mathbf{Q_i}$) with this program by assessing the Pearson correlation between each column and the canonical vectors (set of $\{cv_1, \dots, cv_L\}$). Columns within $\mathbf{Q_i}$ that show a Pearson correlation ≥ 0.6 with the corresponding $cv_i$ and show an average Pearson correlation ≥ 0.2 with $cv_j$ (where j ≠ i) from other matrices, are selected as positive regulators (negative regulators are selected by clearing the same thresholds with the opposite sign). We define the set of selected columns as regulators of a "perturbation program".

5) Multiple perturbation programs can be returned consecutively by this framework. After removing the correlated perturbations identified in step 4) from $\{\mathbf{Q_1}, \dots, \mathbf{Q}_L\}$, steps 3-4 can be repeated until a desired number of programs is reached or no more correlated columns is found. The first program typically encompasses major regulators of a pathway, while the subsequent programs often involve regulators specific to branches of the pathway.

Having identified groups of correlated regulators, we next aimed to enumerate specific gene sets as their downstream targets. To do this, we extracted the previously selected columns of $\{\mathbf{M_1}, \dots, \mathbf{M}_L\}$ from step (5), and merged them into a new matrix $\mathbf{T} \in \mathbb{R}^{G \times S}$, with $S$ representing the number of selected regulators. We performed an eigen-decomposition of **T** and extracted the first eigenvector (of length $G$), which represents an association of each downstream gene with the perturbation program. To establish a baseline, we shuffled the values in each of **T**'s columns individually, and reapplied PCA to generate a 'null' eigenvector. This step was repeated for 200 times to pool all the null eigenvector entries together. Using this pool as a benchmark, we calculated the proportion of null entries exceeding the original entry of each gene. These proportions were the permutation p-values. Genes with p-values below 0.05 or above 0.95 were selected as signature genes of this program/cluster. These genes were then further classified into up- or down-regulated groups according to their effect direction, with down-regulated genes typically marking the primary "pathway signature" due to their expected up-regulation in response to pathway stimulation.

In addition to MultiCCA analysis, which searches for conserved signals across cell lines, we also applied Minimax hierarchical clustering [10] to each of the $\mathbf{Q_i}$ matrices (from step 2 above) individually, thus identifying cell-line specific perturbation clusters. This analysis was performed using the protoclust R package (version 1.6.4) [10]. We set the clustering height parameter to 0.6 to delineate these clusters. These analyses delineate signals that may not be conserved across cell types, and are reported in Supplementary Table 4.

**Obtaining pathway exclusive gene sets**

For closely related pathways, such as IFNβ and IFNγ, which have significantly overlapped pathway signatures, distinguishing between them in pathway enrichment tests is challenging. To address this, we sought to generate exclusive gene sets for each pathway. We started by removing genes shared between two significantly overlapping pathway gene sets. Then, we further eliminated genes from each set that showed DE p-values ≤ 0.05 in any perturbation from the other pathway's perturbation program. The genes remaining after these steps constituted the exclusive gene sets for each pathway. For IFNγ program 1 (involving all core regulators) and 2 (involving subcomponent IRF1/2), we applied a similar procedure to obtain *IRF1*-associated genes (shared genes between program 1 and 2) and *IRF1*-independent genes (program 1 genes that are not shared with program 2).

**Evaluating pathway gene sets in external stimulation datasets**

*Comparison with MSigDB hallmark collection*

We selected the MSigDB hallmark collection [11] (accessible via https://www.gsea-msigdb.org/gsea/msigdb/human/collections.jsp) as a benchmark for our Perturb-seq gene sets. This collection comprises 50 refined gene sets, each containing genes with coherent expression in a specific signaling pathway or biological state. We used the gene sets for IFNβ (INTERFERON_ALPHA_RESPONSE), IFN$\gamma$ (INTERFERON_GAMMA_RESPONSE), TNFα (TNFA_SIGNALING_VIA_NFKB), and TGFβ (TGF_BETA_SIGNALING) from the hallmark collection for comparison.

We assessed our Perturb-seq gene sets using four different external scRNA-seq datasets [12–14], all processed with similar quality control and pre-processing via Seurat. Cells with low number of genes detected (< 200) or high mitochondrial gene content (> 10%) were removed.

*Validation of perturbation programs*

For the IFN$\beta$-stimulated dataset [12] (downloaded from https://github.com/satijalab/seurat-data), we used the original cell type annotations and conducted DE analysis (Wilcoxon rank sum test implemented in Seurat's FindMarkers()) between stimulated and unstimulated cells for each cell type separately, identifying cell-type-specific DE genes due to IFN$\beta$ stimulation. In the IFN$\gamma$ dataset [13] (available via the Gene Expression Omnibus with GEO number GSE178429), since the cell type annotation was unavailable, we utilized Azimuth's human PBMC reference for annotation [4]. Cells with a mapping score below 0.5 were excluded. DE analysis was performed similarly to identify cell-type-specific DE genes. For TNFα and TGFβ-stimulated datasets [14] (available via GEO: GSE147405) involving only one cell line, DE analyses were done between cells at specific treatment time point and non-stimulated cells, yielding time-point-specific DE genes.

After obtaining the significant DE genes (p-value $\leq 0.01$ after Bonferroni correction and log fold change $\geq 0.2$) for each stimulation, we used these DE genes for pathway enrichment analysis based on our Perturb-seq pathway gene sets. Fisher's exact test was used to determine whether the DE genes showed significant overlap with any pathway gene set. We also adopted EASE Score correction to avoid inflation from small DE gene numbers [15]. We also conducted enrichment analysis with pathway exclusive gene sets to differentiate closely related pathways. The same analyses were repeated using MSigDB gene sets.

We validated our IRF1-associated and IRF1-independent gene sets using two external bulk RNA-seq datasets: one from a study on IRF1-deficient patients [16] and another from a study on IRF1 knock-out bat cell lines [17]. We downloaded the raw count matrices for both datasets from the Gene Expression Omnibus (available via GEO: GSE218033 and GSE129390) and imported them into R using the edgeR package (version 3.40.2) [18]. We applied trimmed mean of m-values (TMM) normalization to each sample to adjust for library size differences. After normalization, we converted the count matrices into Seurat objects. To assess the overall activity of our IRF1-associated and -independent gene sets, we conducted module score analyses using the UCell package (version 2.2.0) [19].

*Covid-19 multi-omics blood atlas dataset*

We downloaded the Covid-19 multi-omics blood atlas (COMBAT) scRNA-seq dataset [20] from the Chan Zuckerberg Cell by Gene data portal [21] (https://cellxgene.cziscience.com/collections/8f126edf-5405-4731-8374-b5ce11f53e82), which included cell type annotations. We removed cells with low number of genes detected (< 200) or high mitochondrial gene content (> 10%). We then used Seurat's `AggregateExpression()` function to generate pseudo-bulk data for each cell type within each individual.

To obtain DE results of different disease groups, we followed the original study's workflow [20], applying TMM normalization for each individual using the edgeR package. We then performed pseudo-bulk level DE analyses between patients in specific disease groups and healthy controls using the `glmFit()` function in edgeR. Significant DE genes (p-value $\leq 0.01$ after Benjamini-Hochberg correction and log fold change $\geq 1$) were identified for each cell type and disease group. These genes were used in enrichment analyses using our Perturb-seq pathway gene sets and exclusive pathway gene sets. Additionally, we performed module score analyses for each pathway gene set using the UCell package.

*Crohn's disease dataset*

We downloaded the Crohn's disease scRNA-seq dataset [22] from the Chan Zuckerberg Cell by Gene data portal [21] (https://cellxgene.cziscience.com/collections/5c868b6f-62c5-4532-9d7f-a346ad4b50a7). This data was in Seurat format with cell type annotations. We removed cells with low number of genes detected (< 200) or high mitochondrial gene content (> 10%). We then obtained the DE results for Crohn's disease from the original study (https://ars.els-cdn.com/content/image/1-s2.0-S1074761323000122-mmc4.xlsx). We used the significant DE genes (p-value $\leq 0.05$ after Benjamini-Hochberg correction, log fold change $\geq 0$) in our enrichment analyses, incorporating both our Perturb-seq pathway gene sets and our exclusive pathway gene sets.

*Mouse healing intestine Visium dataset*

We downloaded the mouse healing intestine spatial transcriptomics (Visium) dataset [23] from the Gene Expression Omnibus (GEO number: GSE169749). We first followed the pipeline described in the original study to perform data pre-processing and Harmony integration [24] to integrate the Day 14 (7 days of dextran sodium sulfate administration followed by 7 days of healing) and Day 0 (healthy) samples. We then performed standard clustering analysis based on the mRNA expression data for each spot using Seurat's FindCluster() function. Next, for each cluster, we performed the DE test between the healthy spots and the healing spots to find significant DE genes. These DE genes were then used for pathway enrichment analyses using the pathway gene sets identified in our study.

We also evaluated the pathway activity in the healing tissue by calculating the pathway "induction score" for each spot. We calculated the module scores of each pathway in each spot using the UCell package. For each spot in the healing tissue, we first identified its 20 nearest neighboring spots in the healthy tissue using Seurat's CalcPerturbSig() function. Then, we subtracted the average module score of the 20 spots from the spot's original score. The subtracted score is defined as the pathway induction score for each spot in the healing tissue, and controls for heterogeneity in the healthy tissue (this strategy was inspired by our previous Mixscape study, see [5]) (Figure 6d). We also repeated this analysis using an external TGFβ gene set from the PROGENy database [25], which was used by the original study of the mouse Visium dataset (Figure 6e). We display this score on each spot of the healing tissue, but also on a 'digitally unrolled' version of the mouse colon which restores the original proximal/distal information from the original tissue (we followed the 'unrolling' procedure provided by the authors of the original study at https://github.com/ludvigla/healing_intestine_analysis) (Figure 6f).

**References for Supplementary Methods**

1. Morris, J. A. *et al.* Discovery of target genes and pathways at GWAS loci by pooled single-cell CRISPR screens. *Science* **380**, eadh7699 (2023).

2. Sanson, K. R. *et al.* Optimized libraries for CRISPR-Cas9 genetic screens with multiple modalities. *Nat. Commun.* **9**, 5416 (2018).

3. Simmons, S. K. *et al.* Mostly natural sequencing-by-synthesis for scRNA-seq using Ultima sequencing. *Nat. Biotechnol.* **41**, 204–211 (2023).

4. Hao, Y. *et al.* Integrated analysis of multimodal single-cell data. *Cell* **184**, 3573–3587.e29 (2021).

5. Papalexi, E. *et al.* Characterizing the molecular regulation of inhibitory immune checkpoints with multimodal single-cell screens. *Nat. Genet.* **53**, 322–331 (2021).

6. Ahlmann-Eltze, C. & Huber, W. glmGamPoi: fitting Gamma-Poisson generalized linear models on single cell count data. *Bioinformatics* **36**, 5701–5702 (2021).

7. Mann, H. B. & Whitney, D. R. On a Test of Whether one of Two Random Variables is Stochastically Larger than the Other. *Ann. Math. Stat.* **18**, 50–60 (1947).

8. Jerby-Arnon, L. & Regev, A. DIALOGUE maps multicellular programs in tissue from single-cell or spatial transcriptomics data. *Nat. Biotechnol.* **40**, 1467–1477 (2022).

9. Witten, D. M., Tibshirani, R. & Hastie, T. A penalized matrix decomposition, with applications to sparse principal components and canonical correlation analysis. *Biostatistics* **10**, 515–534 (2009).

10. Bien, J. & Tibshirani, R. Hierarchical Clustering With Prototypes via Minimax Linkage. *J. Am. Stat. Assoc.* **106**, 1075–1084 (2011).

11. Liberzon, A. *et al.* The Molecular Signatures Database (MSigDB) hallmark gene set

collection. *Cell Syst* **1**, 417–425 (2015).

12. Kang, H. M. *et al.* Multiplexed droplet single-cell RNA-sequencing using natural genetic variation. *Nat. Biotechnol.* **36**, 89–94 (2018).

13. Kartha, V. K. *et al.* Functional inference of gene regulation using single-cell multi-omics. *Cell Genom* **2**, (2022).

14. Cook, D. P. & Vanderhyden, B. C. Context specificity of the EMT transcriptional response. *Nat. Commun.* **11**, 2142 (2020).

15. Hosack, D. A., Dennis, G., Jr, Sherman, B. T., Lane, H. C. & Lempicki, R. A. Identifying biological themes within lists of genes with EASE. *Genome Biol.* **4**, R70 (2003).

16. Rosain, J. *et al.* Human IRF1 governs macrophagic IFN-γ immunity to mycobacteria. *Cell* **186**, 621–645.e33 (2023).

17. Irving, A. T. *et al.* Interferon Regulatory Factors IRF1 and IRF7 Directly Regulate Gene Expression in Bats in Response to Viral Infection. *Cell Rep.* **33**, 108345 (2020).

18. Robinson, M. D., McCarthy, D. J. & Smyth, G. K. edgeR: a Bioconductor package for differential expression analysis of digital gene expression data. *Bioinformatics* **26**, 139–140 (2010).

19. Andreatta, M. & Carmona, S. J. UCell: Robust and scalable single-cell gene signature scoring. *Comput. Struct. Biotechnol. J.* **19**, 3796–3798 (2021).

20. COvid-19 Multi-omics Blood ATlas (COMBAT) Consortium. Electronic address: & COvid-19 Multi-omics Blood ATlas (COMBAT) Consortium. A blood atlas of COVID-19 defines hallmarks of disease severity and specificity. *Cell* **185**, 916–938.e58 (2022).

21. CZI Single-Cell Biology Program *et al.* CZ CELL×GENE Discover: A single-cell data

platform for scalable exploration, analysis and modeling of aggregated data. *bioRxiv* 2023.10.30.563174 (2023) doi:10.1101/2023.10.30.563174.

22. Kong, L. *et al.* The landscape of immune dysregulation in Crohn's disease revealed through single-cell transcriptomic profiling in the ileum and colon. *Immunity* **56**, 444–458.e5 (2023).
23. Parigi, S. M. *et al.* The spatial transcriptomic landscape of the healing mouse intestine following damage. *Nat. Commun.* **13**, 828 (2022).
24. Korsunsky, I. *et al.* Fast, sensitive and accurate integration of single-cell data with Harmony. *Nat. Methods* **16**, 1289–1296 (2019).
25. Schubert, M. *et al.* Perturbation-response genes reveal signaling footprints in cancer gene expression. *Nat. Commun.* **9**, 20 (2018).
