## Supplementary Table and File for "Systematic reconstruction of molecular pathway signatures using scalable single-cell perturbation screens": SFile_1_Parse_guide_capture_protocol.pdf

January 29, 2024

### Parse Guide Capture Protocol

- Proceed with the Parse Evercode Whole Transcriptome kit protocol until before cDNA amplification

**At cDNA amplification step:**

- Add guide additive PCR primer (10 µM): 1 µl in each 100 µl sublibrary
- Use the following cycling conditions (modified from the original Parse protocol):

| Step | Cycles |
|---|---|
| 95°C for 3 min | |
| 98°C for 20 sec | 10-18 cycles* |
| **65°C for 45 sec** | 10-18 cycles* |
| 72°C for 3 min | 10-18 cycles* |
| 72°C for 5 min | |
| 4°C hold | |

* Number of cycles varies depending on the number of input cells, see Parse protocol

**Post-amplification SPRI clean-up:**

- Place the tubes with amplified cDNA against a magnetic rack (high setting) and wait for all the beads to bind to the magnet (~2 min: liquid should be clear). CRITICAL! Do NOT discard the supernatant.
- Transfer 90 µl of the clear supernatant into new 100 µl PCR tubes. Discard the original tubes with magnetic beads.
- Add 90 µl of SPRI beads to each sublibrary. (1X cleanup, modified from the original Parse protocol to preserve shorter random hexamer guide products.)
- Pipette to mix 10 times with the pipette volume set to 150 µl.
- Incubate at room temperature for 5 minutes.
- Place the tubes with amplified cDNA against a magnetic rack (high setting) and wait for all the beads to bind to the magnet (~2 min: liquid should be clear).
- With SPRI beads still against the magnetic rack, aspirate and discard the clear supernatant.
- Without resuspending beads, add 180 µl of 85% ethanol and wait for 1 minute.
- Aspirate and discard the ethanol.
- Repeat the ethanol wash once.
- Remove tubes from the magnetic rack and centrifuge briefly (2-3 seconds).
- Place the tubes back on the rack (low setting) and use a P20 pipette to aspirate any residual ethanol.
- Air dry the beads (2 min) CRITICAL! Do NOT over-dry the beads. Over-drying of beads can lead to substantial losses in yield. "Cracking" of the beads is a sign of over-drying.
- Resuspend beads from each tube in 20 µl (for Mini and Regular kits) or 25 µl (for Mega kits) of molecular biology grade water
- Incubate the tubes at 37°C for 10 minutes to maximize elution of amplified cDNA.
- Place the tubes against the magnetic rack (low setting) and wait for all the beads to bind to the magnet (~2 min: liquid should be clear).

- Transfer 20 or 25 µl of the elutant into new PCR tubes with a P20 pipette. Discard the tubes with SPRI beads. The amplified cDNA (and guide) library is now ready to be quanitified.
- Measure concentration of the library using the Qubit dsDNA HS protocol.
- Run libraries on a 2% gel, BioAnalyzer, or tapestation to check if the expected size distribution is there.

**Preparing cDNA libraries for sequencing:**

- Follow the Parse protocol for fragmentation, adaptor ligation, and sample indexing.

**Guide library preparation (gd PCR1):**

- For every amplified cDNA library setup the following gd PCR1 reaction:

| | **1X** |
|---|---|
| Amplified cDNA | 20 ng |
| 2X KAPA Hifi PCR Master Mix (Kapa Biosystems 07958935001) | 25 µl |
| Guide nested primer (10 µM) | 1.25 µl |
| Guide index primer (10 µM) | 1.25 µl |
| Water | Up to 50 µl |

- Run the following PCR program:

98°C for 3 min
98°C for 20 sec | 20-25 cycles*
60°C for 30 sec |
72°C for 30 sec |
72°C for 5 min
4°C hold

* Number of cycles varies depending on the number of input cells

- Expected product size: ~530 bp

**Post gd PCR1 SPRI clean up:**

- Add 50 µl of SPRI beads to each guide library (1X cleanup) for a total volume of 100 µl.
- Mix well by pipetting 10 times with the P200 pipette volume set to 75 µl.
- Incubate at room temperature for 5 minutes.
- Place the tubes with amplified guides against a magnetic rack (high setting) and wait for all the beads to bind to the magnet (-2 min: liquid should be clear).
- With SPRI beads still against the magnetic rack, aspirate and discard the clear supernatant with a pipette.
- Without resuspending beads, add 180 µl of 85% ethanol and wait for 1 minute.
- Using a pipette, aspirate and discard the ethanol from each tube.
- Without resuspending beads, add another 180 µl of 85% ethanol and wait for 1 minute.

- Using a pipette, aspirate and discard the ethanol from each tube.
- Remove tubes from the magnetic rack and centrifuge quickly (2-3 seconds).
- Place tubes back on the rack (low setting) and with a P20 pipette remove any residual ethanol.
- With the tubes still on the rack, air dry the beads (~2 min). CRITICAL! Do NOT over-dry the beads. Over-drying of beads can lead to substantial losses in yield. "Cracking" of the beads is a sign of over-drying.
- Remove tubes from the magnet and resuspend beads in 20 µl of molecular biology grade water.
- Incubate at room temperature for 5 minutes.
- Place the tubes against the magnetic rack (low setting) and wait for all the beads to bind to the magnet (~2 min: liquid should be clear).
- Transfer 20 µl of the elutant into new PCR tubes with a P20 pipette. Discard the tubes with SPRI beads. The amplified guide library is now ready to be quantified.
- Measure concentration of the cDNA using the Oubit dsDNA HS protocol.
- Run libraries on 2% gel, BioAnalyzer or tapestation to check if the expected size distribution is there.

**Guide library preparation (gd PCR2):**

- For every amplified cDNA library setup the following gd PCR2 reaction:

| | **1X** |
|---|---|
| gd PCR1 product | 20 ng |
| 2X KAPA Hifi PCR Master Mix (Kapa Biosystems 07958935001) | 25 µl |
| P7 primer (10 µM) | 1.25 µl |
| SI-PCR primer (10 µM) | 1.25 µl |
| Water | Up to 50 µl |

- Run the following PCR program:

98°C for 3 min
98°C for 20 sec
60°C for 30 sec
72°C for 30 sec
(98°C for 20 sec – 72°C for 30 sec: 5-7 cycles*)
72°C for 5 min
4°C hold

* Number of cycles varies depending on the number of input cells

- Expected product size: ~575 bp

**Post gd PCR2 SPRI clean up:**

- Add 50 µl of SPRI beads to each guide library (1X cleanup) for a total volume of 100 µl.
- Mix well by pipetting 10 times with the P200 pipette volume set to 75 µl.
- Incubate at room temperature for 5 minutes.

- Place the tubes with amplified guides against a magnetic rack (high setting) and wait for all the beads to bind to the magnet (-2 min: liquid should be clear).
- With SPRI beads still against the magnetic rack, aspirate and discard the clear supernatant with a pipette.
- Without resuspending beads, add 180 µl of 85% ethanol and wait for 1 minute.
- Using a pipette, aspirate and discard the ethanol from each tube.
- Without resuspending beads, add another 180 µl of 85% ethanol and wait for 1 minute.
- Using a pipette, aspirate and discard the ethanol from each tube.
- Remove tubes from the magnetic rack and centrifuge quickly (2-3 seconds).
- Place tubes back on the rack (low setting) and with a P20 pipette remove any residual ethanol.
- With the tubes still on the rack, air dry the beads (~2 min). CRITICAL! Do NOT over-dry the beads. Over-drying of beads can lead to substantial losses in yield. "Cracking" of the beads is a sign of over-drying.
- Remove tubes from the magnet and resuspend beads in 20 µl of molecular biology grade water.
- Incubate at room temperature for 5 minutes.
- Place the tubes against the magnetic rack (low setting) and wait for all the beads to bind to the magnet (~2 min: liquid should be clear).
- Transfer 20 µl of the elutant into new PCR tubes with a P20 pipette. Discard the tubes with SPRI beads. The amplified guide library is now ready to be quantified.
- Measure concentration of the cDNA using the Oubit dsDNA HS protocol.
- Run libraries on 2% gel, BioAnalyzer or tapestation to check if the expected size distribution is there.

**Oligonucleotide sequences (5' -> 3'):**

- Guide additive PCR primer:
  - GAGGGCCTATTTCCCATGATT*C*C
- Guide nested PCR primer
  - CTACACGACGCTCTTCCGATCTGTGGAAAGGACGAAACACC
- Guide index primer (P7-i7-R2):
  - CAAGCAGAAGACGGCATACGAGATATGTCCGTGTGACTGGAGTTCAGACGTGTGC
  - Use a different i7 index for each sublibrary
- 10X Genomics SI-PCR primer:
  - AATGATACGGCGACCACCGAGATCTACACTCTTTCCCTACACGACGC*T*C
- P7 primer:
  - CAAGCAGAAGACGGCATACGA
